# Atomic-Level Investigation of *KCNJ2* Mutations Associated with Ventricular Arrhythmic Syndrome Phenotypes

**DOI:** 10.1101/2024.10.01.616187

**Authors:** Saba Munawar, Corey L Anderson, Louise Reilly, Ryan Woltz, Yusra Sajid Kayani, Nipavan Chiamvimonvat, Lee L. Eckhardt

## Abstract

*KCNJ2* encodes the inward rectifying potassium channel (Kir2.1) that creates *I_K1_* which maintains the cardiac resting membrane potential and regulates excitability. Mutations in *KCNJ2* have been linked to several clinical phenotypes associated with ventricular arrhythmia and sudden death including Andersen-Tawil syndrome (ATS) related to loss of function mutations and Short QT Syndrome 3 related to gain of function mutations. Detailed structural-functional relationships to explain the arrhythmia phenotypes are understudied and limit the capacity to provide precision medicine. Here, we combine in-depth and complementary computational molecular modeling techniques with functional analysis from three patients with ATS that harbor *KCNJ2* mutations R67Q, R218L, and G300D. Whole-cell patch-clamp experiments revealed loss of function in homomeric mutant channels. Full-length Kir2.1 models were developed for structure-based investigation, and mutations were introduced in both open and closed conformations. Site-directed mutagenesis identified altered interaction profiles contributing to structural perturbations. Molecular dynamics simulations assessed the impact of each mutation on overall channel conformation and stability. Principal component analysis and normal mode analysis revealed mutation-specific structural perturbations. These findings extend beyond previous studies, offering atomic-level characterization of mutation-specific perturbations. Our multifaceted approach provides first atomic-level insights into the molecular mechanisms underlying ATS, paving the way for targeted therapeutic strategies.

**Study Highlights:** 1) Clinical mutation analysis confirmed loss-of-function.

2) Mutation analysis revealed that these clinical mutations dramatically alter the interaction pattern of the mutated residue and subsequently disturbs channel stability.

3) The molecular dynamics based RMSD and RMSF evaluations show that the open conformation state of the channel is more stable comparative to the closed state however, the mutations impact channel conformations regardless of conductance state.

4) The PCA (principal component analysis) and PCA based NM (normal mode) analysis revealed that these clinical *KCNJ2* mutations caused significant conformational changes, even distant from the specific residue.

4) This study is the first extensive *in silico* and experimental analysis of *KCNJ2* clinical mutations that start from the arrhythmia phenotype and lead to an in-depth atomic-level investigation. These newly resolved features pave the way towards a better understanding of the molecular disease mechanism and new therapeutic strategies.

## Introduction

*KCNJ2* encodes inward rectifier potassium channel 2.1 (Kir2.1), which forms the dominant channel which conducts *I_K1_*. Loss-of-function mutations are associated with Andersen-Tawil syndrome type 1 (ATS1), while the gain-of-function mutations are associated with short QT syndrome 3 and familial atrial fibrillation^1^. Some loss-of-function mutations have arrhythmia phenotype that is dependent on adrenergic stimulation^2,3^. We and others have characterized functional effects of loss-of-function (LOF) mutations in both cellular and transgenic animal models^3–5^. However, less is known about the specific Kir2.1 molecular and atomic structural perturbations that cause channel dysfunction. This is in part due to the lack of available open and closed high-resolution Kir2.1 channel structure. Recently, a cryo-EM structure of Kir2.1 in the closed conformation state with a resolution of 4.30 Å was reported^6^, and a high-resolution crystal structure of Kir2.2 was previously published^7^. Both structures omit the distal N- and C-termini of the channel, which limits the applications of these models for comprehensive *KCNJ2* mutational analysis and overall conformational change.

To better resolve the structural and functional consequences of clinical *KCNJ2* mutations, we created and functionally assessed the impact of three mutations from the atomic level to the 3D tetrameric structure followed by the assessment of ion channel current. We developed the first full-length open conformational state homology model by taking advantage of the Kir2.2 crystal structure (PDB ID: 6M84)^8^ that shares a close homology with Kir2.1. To reduce computational time, we introduced the mutations in both open and closed conformation structures individually and compared them with the native Kir2.1-WT structure to study the complete behavior of mutation change.

ATS mutations under investigation have been previously reported and correlate to ATS patients from the University of Wisconsin Inherited Arrhythmia Clinic and functionally demonstrate loss of function^2–4^. Structurally, the R67 and R218 residues are located near the slide helix, where regulatory ligands bind^9^. However, G300 is part of the G-loop region that forms a girdle around the central pore and creates a diffusion barrier between the transmembrane domain and cytoplasmic pore^10^. G-loop is a hotspot for ATS-causing mutations that alter the cytoplasmic K^+^ conduction pathway and lower gate function^11–14^, although its precise role in channel gating remains unresolved. Therefore, we focused on three ATS mutations to understand how these specific mutations cause conformational changes and structural instability that alters the normal channel behavior. We used a comprehensive approach combining functional and structural characterization techniques, including cellular electrophysiology, molecular modeling, and computational simulation. These analyses provide deeper insights into how specific mutations alter interaction profiles, leading to overall structural conformational changes. Our approach, by combining experimental and computational methods, offers a detailed understanding of the structural and functional consequences of ATS mutations in the Kir2.1 channel. To our knowledge, this study is the first multifaceted *in silico* and *in vitro* analysis of *KCNJ2* clinical mutations that start from the arrhythmia phenotype and lead to an atomic-level investigation. These findings contribute to our knowledge of channelopathies and may inform future therapeutic strategies for ATS.

## Results

### Functional characterization of *KCNJ2* mutations

Clinical mutations (R67Q, R218L and G300D) were chosen from patients in the UW Inherited Arrhythmia clinic and have been reported elsewhere^2,3^. Recordings from the stable cell lines are shown in Fig. 1. Representative traces (Fig. 1a) and the current-voltage (I/V) plot (Fig. 1b) show the typical “N-shaped” inward rectifier current for Kir2.1-WT with a reversal potential close to Keq. Homo-tetrameric R67Q, R218L and G300D Kir2.1 channels lack meaningful current (Fig. 1a and Fig. 1b). Quantitative analysis of outward *I_K1_* at −50 mV revealed significant differences between Kir2.1-WT and mutant channels (Fig. 1c). Kir2.1-WT demonstrated robust outward current (1.735 ± 0.389 pA/pF, n=8), while R67Q (−0.3652 ± 0.389 pA/pF, n=7), R218L (−0.1253 ± 0.389 pA/pF, n=8), and G300D (−0.01909 ± 0.389 pA/pF, n=11) showed negligible or slightly negative currents (p < 0.001 for all comparisons). Western blot analysis all mutant constructs express (Fig. 1d), and run at the predicted size.

**Fig. 1.**
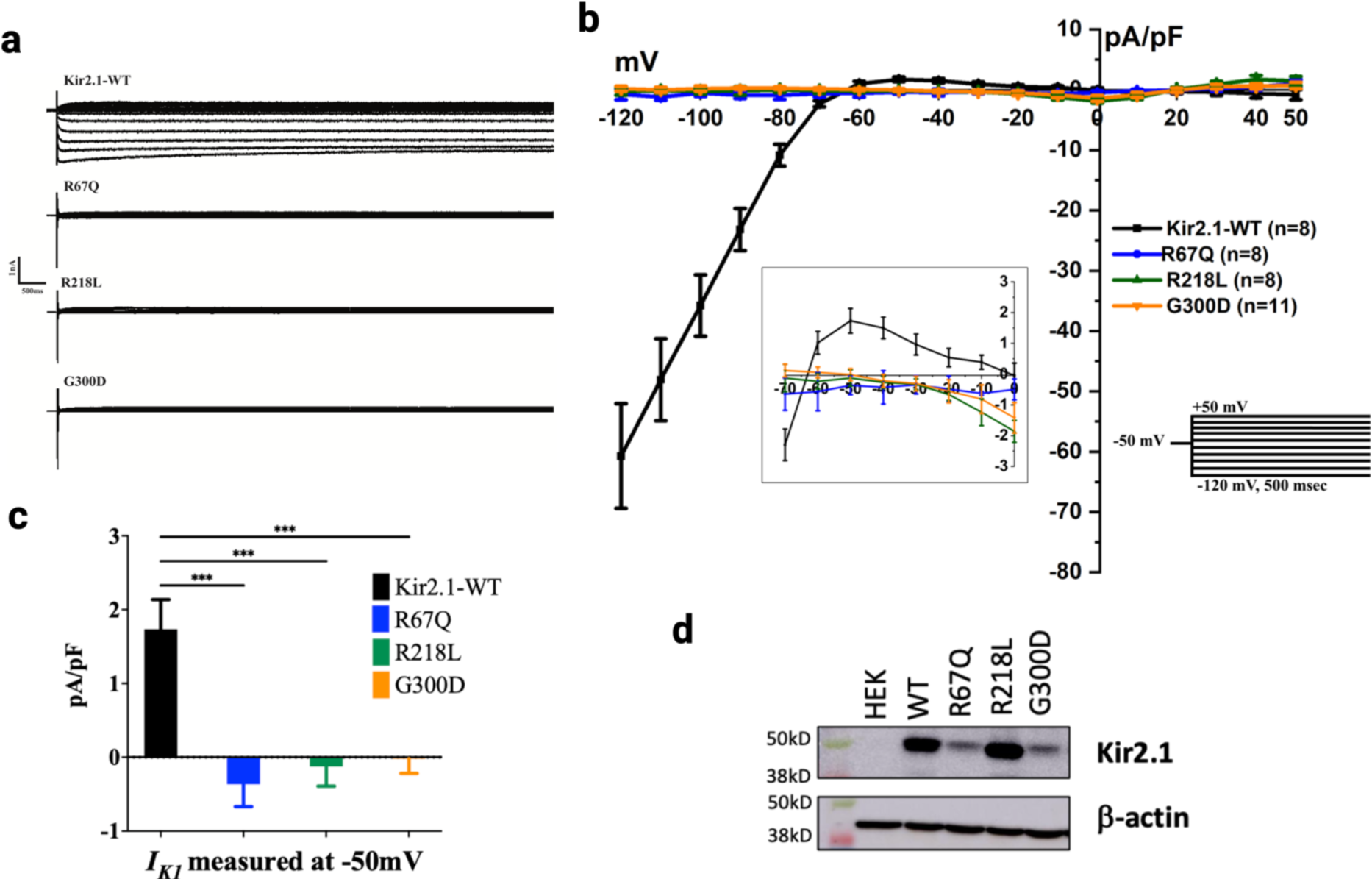
Functional assessment of Kir2.1-WT and clinical mutation. **a)** Representative whole cell current traces are obtained by stepping the membrane from a holding of −50mV to voltages ranging from −120 to +50mV. **b)** Mean Steady-state current-voltage relationship for WT Kir2.1 with inset highlighting the outward current at physiologic voltages. Currents are calculated following barium subtraction. The error bar represents the standard error of the mean (SEM) **c)** Mean outward *IK1* current measured at −50mV, and all three mutants showing no current at baseline compared to WT. Data expressed as mean ± SEM. Mutants show significantly reduced current compared to WT. Statistical significance determined by one-way ANOVA with Dunnett’s multiple comparisons test (*** p < 0.001 for all mutants vs. WT) **d)** Western blot of lysates from HEK stable cell lines expressing each of the clinical Kir2.1 mutations along with ß –actin loading.

### Structure modeling of Kir2.1 and selected clinical mutations

To determine the mechanism of a single amino acid change that leads to loss of function, we performed in-depth molecular modeling. We used the recently published cryoEM structure (PDB ID: 7ZDZ^6^) for the closed conformation state with ∼40 and ∼60 missing residues from N and C-terminal regions, respectively. However, due to the lack of an experimental human open-state Kir2.1 structure, we constructed a full length homology model based on the crystal structure of Kir2.2 (PDB ID: 6M84^8^) with a resolution of 2.81Å (77% query coverage, 75.60% sequence identity). 100 independent models were developed using MOE v2022^15^, whereby model refinement and scoring were performed through GB/VI functions. The top five models with GB/VI scores in the range −7.77 x104 to −7.34 x104 were further assessed using ERRAT (score: 92.1-92.4) and PROCHECK. The model with optimal validation parameters was selected as the final model for further computational studies. The final model for the open-state Kir2.1 channel (Fig. 2a) exhibited the best model validation parameters i.e., ERRAT scores of 92.4, and the Ramachandran analysis revealed 98.6% of residues in the favorable regions (Supplementary Fig.1). The van der Waals surface of ion conduction pore was generated using HOLE software^16^, which revealed the upper/ helix bundle crossing (HBC) and lower or G-loop gating points (Fig. 2b). The closed state cryoEM structure^6^ revealed that I176 and M180 are key residues involved in HBC or upper gating region and A306 is important for cytoplasmic or lower gating region. However, our in-house open conformation state model, showed that the G177 and M307 residues are also important in upper and lower gating, respectively. The plot of the pore radius between van der Waals surfaces of open and closed conformation highlighting the gating residues is provided in <u>Supplementary</u> Fig. 1 <u>and</u> Supplementary Fig. 2. The selectivity filter, upper gating residues and lower gating residues that are also part the of G-loop region are highlighted in Fig. 2c. Furthermore, mutations under investigation are individually introduced to the channel in the open and closed conformation. Fig. 2d shows the location of specific mutation and inset indicates the amino acid change and their respective functional group.

**Fig. 2.**
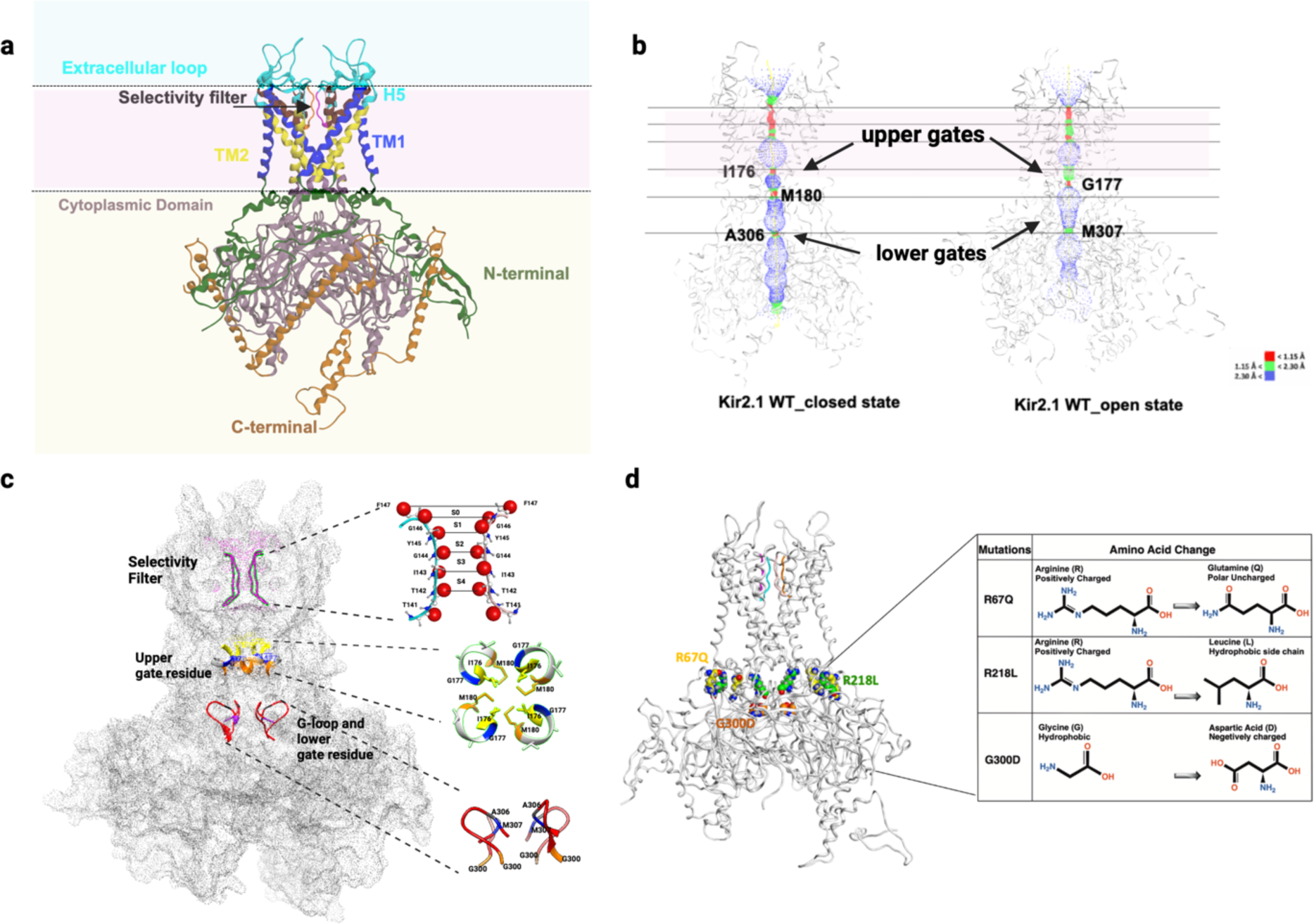
Kir2.1 full-length 3D structure and pore analysis. **a)** Full-length Kir2.1 structure in open conformation with complete N and C-terminal regions **b)** Pore radius of open and closed conformation also showing gating residues/construction points. Upper gates are indicating the helix bundle crossing region. The lower gate is by the G-loop region **c)** Open channel surface with enhanced structural feature Selectivity filter (SF) using ball-and-stick representation with carbonyl oxygen atoms (red balls). Black lines represent the distances between backbone oxygen of SF residues (141-147). Five distinct positions of K^+^ in the SF, sites S0-S4 are labeled. The upper gate residue inset showing I176, G177 and M180 forms the gating structure. G-loop inset showing the lower gating residues A306 and M307 **d)** Location and amino acid change (inset) for the *KCNJ2* mutations studied in this article.

### Site-directed mutagenesis analysis

MOE v2022.02^17^ was used to systematically introduce the single mutations at appropriate positions in all chains of one structure at a time, considering both the open and closed states of Kir2.1. As a result, three open- and three closed-state Kir2.1 mutant structures were obtained. To elucidate changes in interactions and stability of the structure, mutational analysis were performed with reference to the WT-open and WT-closed structures of Kir2.1. We investigated the interaction pattern of the selected residues in WT and mutant state with other residues present in the vicinity, and the involvement of these residues in backbone stability of the structure. Additionally, the structural overlays of mutant and WT conformations provide crucial insights into bond remodeling and localized structural deviations. These structural perturbations collectively contribute to channel instability, offering a molecular basis for the observed functional deficits.

The R67Q mutation was analyzed in both the open and closed states (Fig. 3) in the homomeric Kir2.1 channel. In the WT-closed state of the channel, a hydrogen bond at 2.18Å was observed between the chain2 residues 2: R67 (for 2: R67, 2 refers to chain number, and R67 is the position of arginine) and 2: T74. Three additional hydrogen bonds were observed between chain3 residues 3: R218 and 3: E191, with distances between 1.75-2.16Å. Another hydrogen bond at 1.81Å was mediated by the residues 2: G65 and 3: R218. Other residues involved in hydrogen bonding are 2: T75, 2: D71, and 2: T74 (Fig. 3a and Supplementary Table 1). Furthermore, in the R67Q-mutant closed structure, the hydrogen bonding pattern depicted by 2: R67 was lost, and two new interactions were observed between the residues 2: Q67-2: D71 and 2: Q67-2: L69, which indicates a displacement upon mutation (Fig. 3a and Fig. 3b). However, in the WT-open state of Kir2.1, the R67 residue was involved in an intricate hydrogen bonding pattern, in which five hydrogen bonds with distances in the range 1.70-3.32 Å were observed. Four of these hydrogen bonds were observed between 1: R67 residue of chain 1 and 3: E191 of chain 3 whereas, one hydrogen bond at 2.63Å was formed between 1: R67 and 1: E63 (Fig. 3c). The dense hydrogen bonding pattern of 1: R67 indicates that it plays an important role in maintaining the structural stability and compactness of the alpha helix. Additionally, three additional hydrogen bonds were observed in the near vicinity of 1: R67 residue between the residues 3: T192-1: D71, 1: E63-1: E63, and 3: L217-1: Y68, respectively (Fig. 3c). The detailed interaction patterns for WT and Q67 mutant structures are presented in Supplementary Table 1. Notably, R67Q mutation in the WT open state replaces the basic amino acid 1: R67 with a polar uncharged 1: Q67, which disrupts the hydrogen bonding pattern. The 1: Q67 residue forms a single hydrogen bond interaction with 1: D71 at a distance of 1.89Å. An additional H-Pi interaction was also observed between chain 1 residues of Kir2.1, namely 1: Y68 and 1: K64. The remaining hydrogen bonds formed in the WT open structure remain intact in the mutant structure with slight deviations in the distances, as shown in Fig. 3d and Supplementary Table 2.

**Fig. 3.**
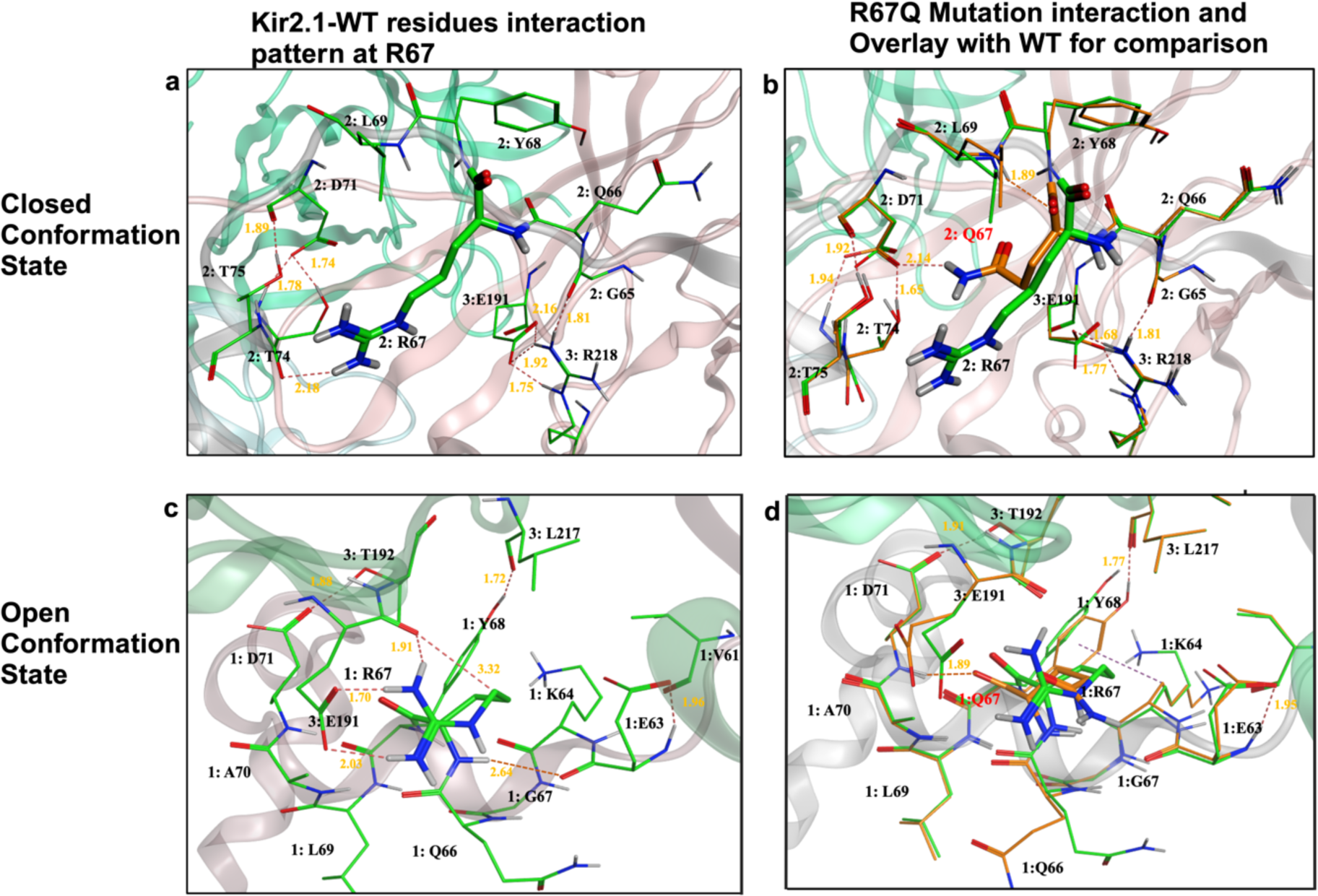
Site-directed mutation analysis of R to Q change at location 67 in open and closed state of Kir2.1. **a)** Interaction profile of R67 with neighboring residues in closed conformation state highlighting the hydrogen bond interaction **b)** Overlay of WT structure and change in residue from R to Q and change in hydrogen bond interactions **c)** Interaction profile of R67 with neighboring residues in open conformation state highlighting the hydrogen bond interaction **d)** Overlay of WT structure and change in residue from R to Q and change in hydrogen bond interactions in open conformation state of the channel.

Similarly, to explore the effect of R218L mutation, the interaction patterns in the WT and R218L-mutant structures were compared. In the Kir2.1-WT open and closed state, R218 from chain 1 was involved in a dense hydrogen bonding pattern. Four hydrogen bonds were mediated by 1: R218 in the closed state with the residues including 1: S220, 1: T192 and 1: N216, as well as between 1: L217 and 4: K64. The detailed interactions are presented in Fig. 4a and Supplementary Table 1. In the closed state, replacement with 1: L218 led to loss of the dense hydrogen bonding pattern of 1: R218. In comparison to the WT-closed state structure, an additional hydrogen bond at 2.18Å was observed between 4: K64 and 4: R67 (Fig. 4b and Supplementary Table 1) upon replacement with 1: L218. In the WT-open state Kir2.1 structure 1: R218 formed four hydrogen bonds with the residues 1: N216, 4: D71 and 4: T75 (distances between 1.80 to 1.94 Å) (Fig 4c). The L218-mutant replacement of the arginine (basic and large residue) with the non-polar smaller residue 1: L218 resulted in the loss of the dense hydrogen bonding pattern previously mediated by 1: R218. In the R218L-mutant open state only one hydrogen bond was observed between the mutated residue 1: L218 and 1: N216 at 1.91Å.

**Fig. 4.**
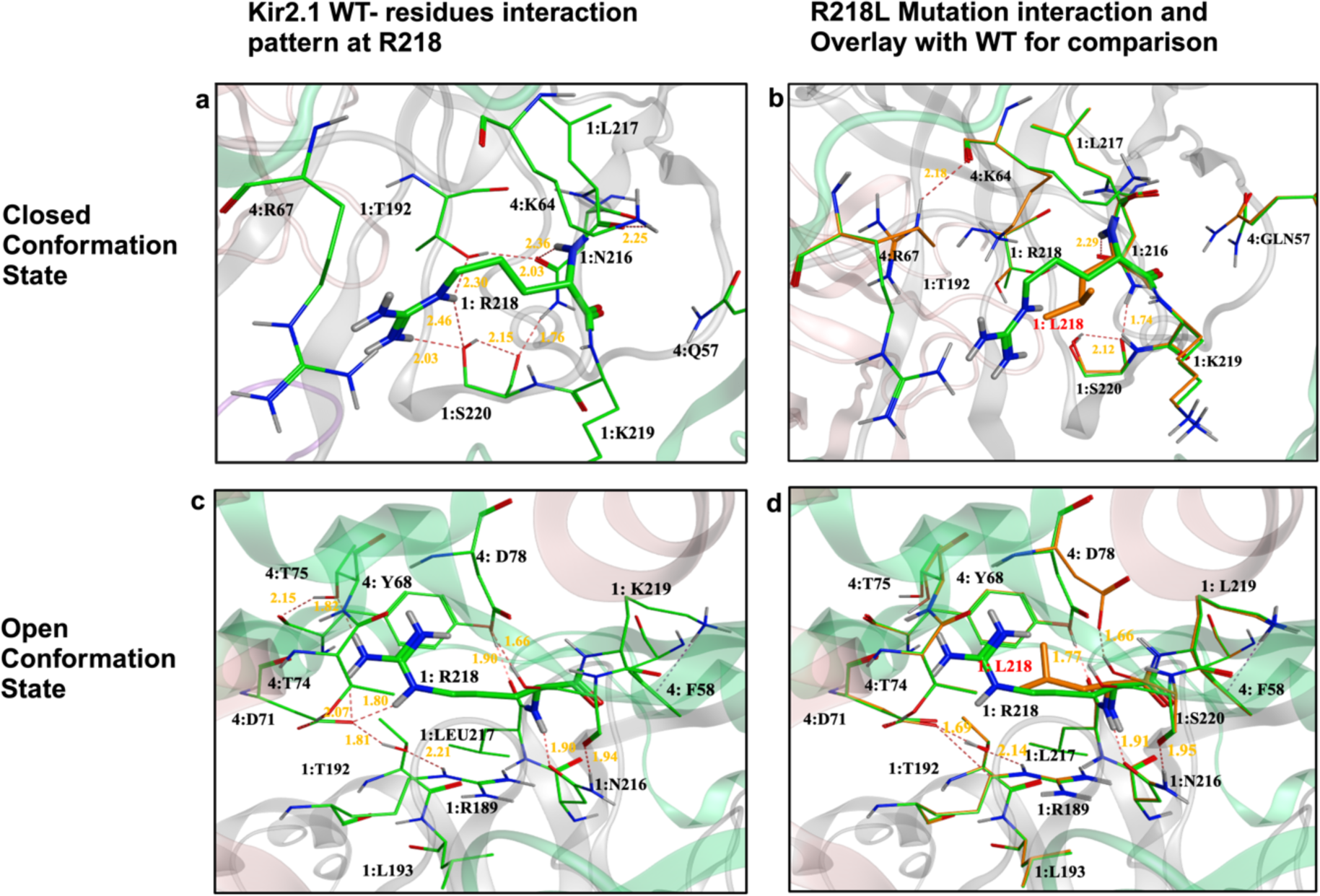
Site-directed mutation analysis of R to L change at location 218 in open and closed state of Kir2.1. **a)** Interaction profile of R at location 218 with the neighboring residues in closed conformation state highlighting the hydrogen bond interaction **b)** Overlay of WT structure and change in residue from R to L and change in hydrogen bond interactions **c)** Interaction profile of R at location 67 with the neighboring residues in open conformation state highlighting the hydrogen bond interaction **d)** Overlay of WT structure and change in residue from arginine to L and change in hydrogen bond interactions in open conformation state of the channel.

However, other hydrogen bonds formed between the residues 1: L217-4: Y68, 1: S220-4: D78,1: S220-1: N216 and 1: T192-4: D71 persisted in the mutant structure with slight changes in distances (Fig.4d and Supplementary Table 1). Overall, the dense network of hydrogen bonds formed by the extended guanidine group of the arginine side chain in the WT open state and the intra hydrogen bond formed by 4: T75 at 2.15 Å is completely lost upon L218 mutation. However, the remaining inter-residue hydrogen bonds present in the near vicinity of the R218 remain intact with modified distances upon substitution with L218.

Mutational analysis was also performed for G300D in both the open and closed state Kir2.1. In the WT-closed state three hydrogen bonds within the distance range of 1.90-2.48Å were observed between 1: G300 and 1: T309. Other hydrogen bonding interactions were observed between the residues 1: M301-1: V302, 1: E224-1: E224, 1: T308-1: T308 (Fig. 5a). Additional information on the detailed interactions is provided in Supplementary Table 1. Notably, in the mutant state, the G300D substitution might not only introduce bulkiness but also add a negative charge, which could possibly lead to new interactions (i.e., electrostatic, hydrogen bonding) with nearby residues or disruption of local structure. This might also lead to the stabilization or the disruption of existing interactions. However, upon replacement with the bulkier residue D300, a hydrogen bond is maintained with the residue 1: T309, however the distance is increased to 3.02Å from 2.48Å that was observed in the WT closed structure with 1: G300. This increase in the distance might be due to the replacement of the smallest amino acid G300 with a bulkier residue D300, which has a larger sidechain in comparison. In addition, another hydrogen bonding interaction between the HG1 of 1: T309 and OD2 of 1: D300 was observed at a distance of 1.62Å. However, the hydrogen bond previously formed in the WT structure between O of 1: T309 and H of 1: G300 was lost upon mutation. Other hydrogen bonds observed in the close vicinity of G300 in the WT closed structure remain intact with slightly modified distances (Fig. 5b). Most prominently G300 is the most critical residue that forms the defining component of the flexible G-loop structure, which is crucial for the inward rectification of the Kir2.1 ion channel^18^. The flexibility and positioning of the G-loop is essential for selective permeability and gating mechanisms. Therefore, the replacement of G300 with the bulkier and less flexible D300 might lead to reduced flexibility and potentially altered conformation.

**Fig. 5.**
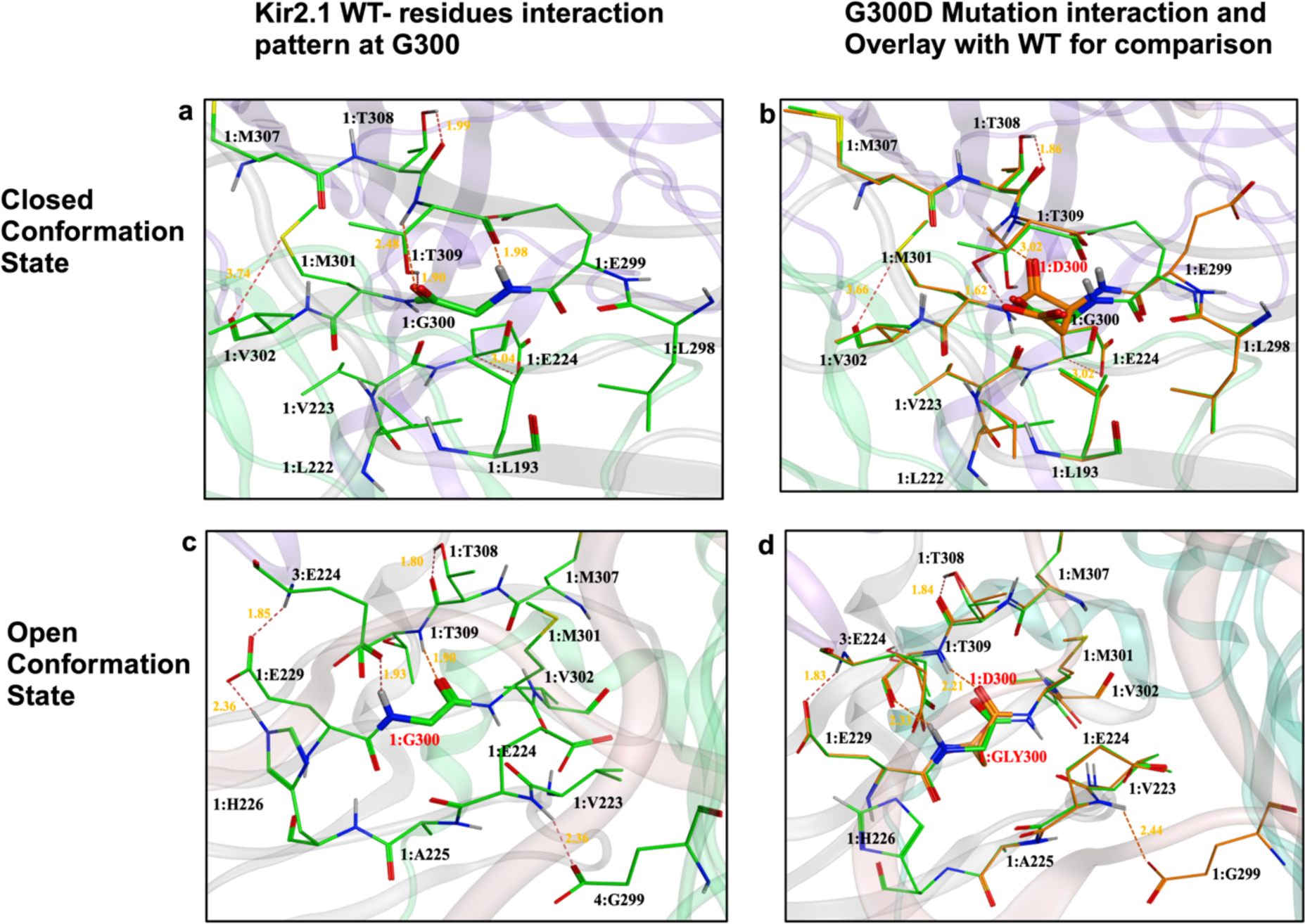
Site-directed mutation analysis of G to D change at location 300 in open and closed state of Kir2.1. **a)** Interaction profile of GLY at location 300 with the neighboring residues in closed conformation state highlighting the hydrogen bond interaction **b)** Overlay of WT structure and change in residue from GLY to ASP and change in hydrogen bond interactions **c)** Interaction profile of GLY at location 300 with the neighboring residues in open conformation state highlighting the hydrogen bond interaction **d)** Overlay of WT structure and change in residue from GLY to ASP and change in hydrogen bond interactions in open conformation state of the **a)** PCA plot explains the projection of molecular dynamics trajectories onto first two principal components (PC1 and PC2). Each point represents a snapshot from the trajectory, and clusters indicate regions of the conformational space where the protein spends significant time, corresponding to energy minima. The color density indicates the population density of the snapshots, with yellow representing higher density and purple representing lower density. for Kir2.1-WT in closed state **b)** PCA plot of R67Q in closed state **c)** PCA plot of R218L in closed state **d)** PCA plot of G300D in closed conformation state. **e)** PCA plot of R67Q **in** open state **f)** PCA plot of R218L in open state **g)** PCA plot of G300D in closed conformation state. The structure underneath each plot is showing the represented frame structure overlay on refence frame in open conformation state.

In the WT-open state Kir2.1, hydrogen bonds were observed between the residues 1: E229-3: E224, 1: E229-1: H226, 1: E224-4: E229, and 1: T308-1: T308 with distances less than 2.5Å. Notably, 1: G300 mediated a hydrogen bonding interaction at 1.93Å with 3: E224. In the WT-open state Kir2.1, hydrogen bonds were observed between the residues 1: E229-3: E224, 1: E229-1: H226, 1: E224-4: E229, and 1: T308-1: T308 with distances less than 2.5Å. Notably, 1: G300 mediated a hydrogen bonding interaction at 1.93Å with 3: E224. In the WT-open state Kir2.1, hydrogen bonds were observed between the residues 1: E229-3: E224, 1: E229-1: H226, 1: E224-4: E229, and 1: T308-1: T308 with distances less than 2.5Å. Notably, 1: G300 mediated a hydrogen bonding interaction at 1.93Å with 3: E224. Another hydrogen bond at 1.90 Å was observed between 1: G300 and 1: T309 (Fig. 5c). The 1: G300-3: E224 hydrogen bond is lost (Fig. 5d) upon replacement with 1: D300. However, upon G300D mutation in the structure, the previously established intermolecular hydrogen bonds in the WT-open state slightly changes the distances as shown in Fig.5d. Two new hydrogen bonds with distances in the range 2.21-2.33Å were observed between 1: D300 and 1: T309 (Fig. 5d and Supplementary Table 1).

For all WT-open, WT-closed, mutant-open and mutant-closed structures a static view is presented, which gives an overview of the effects of a mutation on the Kir2.1 channel in the local area, showing how a single residue mutation changes the interaction profile of all the residues present in the vicinity. To elucidate the global impact of a particular mutation on the interaction profile and structural stability in dynamic environments, molecular dynamics simulations were performed for the systems under investigation.

### Molecular dynamics simulation analysis

To further investigate the effect of specific mutations on structural dynamics, interactions, and conformational changes, a molecular dynamics (MD) simulation for each WT and mutant in open and closed state was performed for 500 ns with an additional 80 ns of extended equilibrium protocol (Supplementary Fig. 3). Highly flanking N-C terminal regions were trimmed to avoid large RMSD and RMSF values at the very ends of the structures. This simulation was conducted using Amber22^19^on the high-performance computing (HPC) cluster Expanse (Full details in Methods section), followed by the determination of C_alpha_ Root Mean Square Deviation (C_α_RMSD) and C_alpha_ Root Mean Square Fluctuation (C_α_RMSF). After MD simulations of Kir2.1, the C_α_RMSDs were calculated for WT and mutant systems in both open and closed states of the channel. The C_α_RMSDs for the proteins (rmsd_protein), selectivity filter (rmsd_SF) and Pore domain (rmsd_PD) were analyzed to assess the stability of the WT and mutant structures of Kir2.1. Notably, for the WT-closed structure of Kir2.1, higher fluctuations up to 9Å were observed in the protein, while the SF and PD appear to be relatively stable as indicated by C_α_RMSD values below 5Å during the 600ns MD simulations (Supplementary Fig. 4). The overall RMSD plot in closed conformation state of protein is shown in Fig. 6b.

**Fig. 6.**
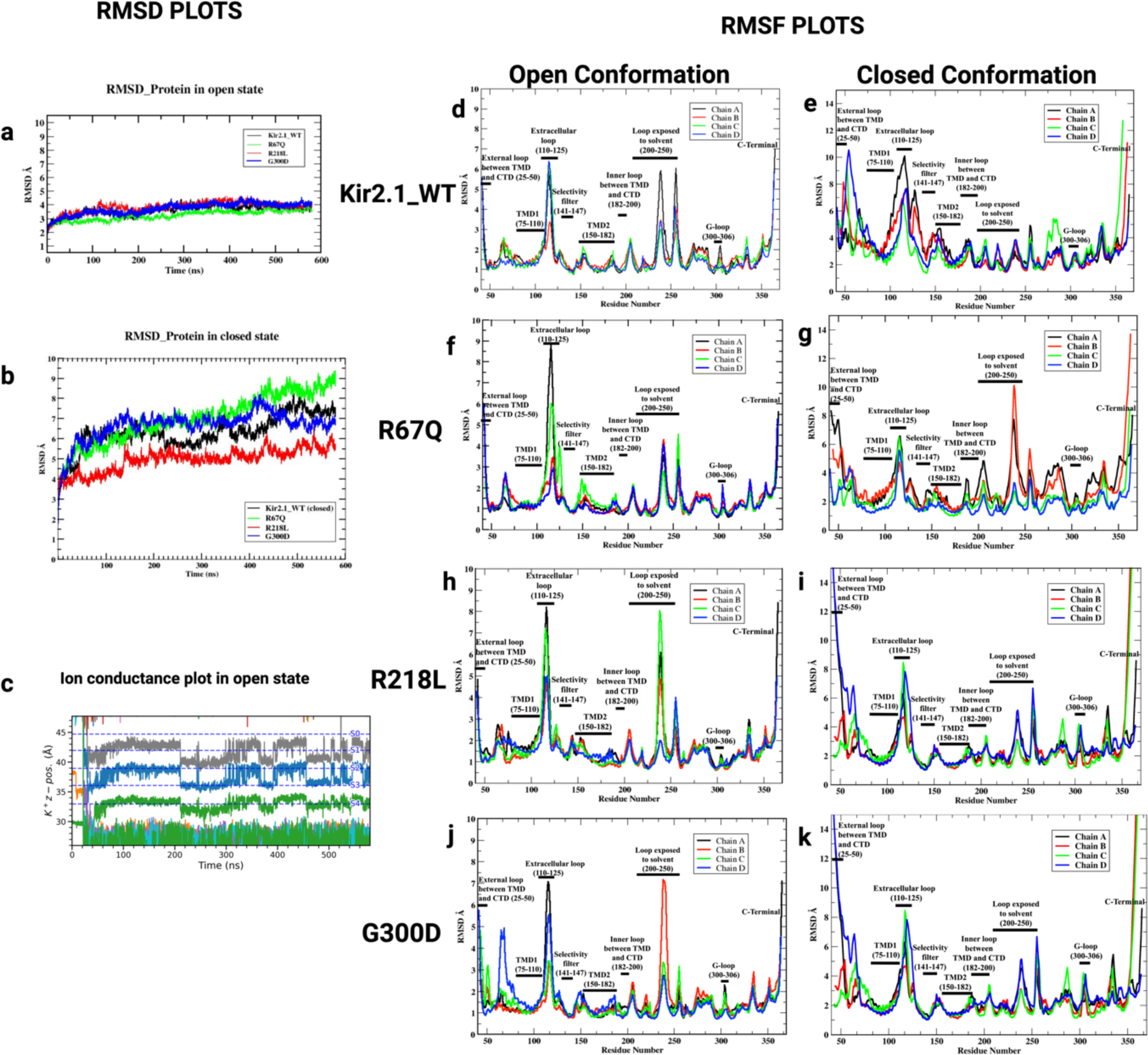
RMSD and RMSF plots of Kir2.1-WT and clinical mutants (R67Q, R218L, G300D) in opened and closed conformation State. **a)** Combined RMSD plot of Kir2.1-WT and mutants overall protein in open conformation state **b)** Combined RMSD plot of Kir2.1-WT and mutants overall protein in closed conformation state and closed state. **c)** showing movement of ion through selectivity filter in open state of the channel during simulation time **d)** RMSF plots for each Kir2.1-WT open state showing all 4 chain the x-axis showing residue number and Y-axis is labelled as RMSD in Å that showing residue fluctuation throughout simulation time from reference structure RMSF plots for each Kir2.1-WT and Mutants showing 4 chain and x-axis are labeled with residue number and Y-axis is labelled as RMSD in Å that showing residue fluctuation throughout simulation time from reference structure. **e)** RMSF plot of Kir2.1-WT in closed state **f)** RMSF plot of R67Q mutation in closed state g) **g)** RMSF plot of R67Q mutation in open state **h)** RMSF plot of R218L mutation in closed state **i)** RMSF plot of R218L mutation in open state **j)** RMSF plot of G300D mutation in closed state **k)** RMSF plot of G300D mutation in open state.

Contrary to the WT-closed structure, the WT-open structure depicts higher stability as shown by the lower C_α_RMSD values (< 4.2 Å) for the protein SF and PD, respectively (Supplementary Fig. 4). This can be attributed to the greater structural variability or conformational changes occurring in the closed state in comparison to the open state of Kir2.1. In addition to the WT-structures, the C_α_RMSDs were also analyzed for the mutant structures. For the R67Q-closed mutant structure a similar pattern (as of the WT-closed structure) was observed in the C_α_RMSDs of SF and PD (C_α_RMSD <4.2 Å), whereas higher C_α_RMSD fluctuations up to 10Å were observed in the protein (Supplementary Fig. 4). However, in the R218L-closed mutant structure lower C_α_RMSD values of <7Å, <4 Å and <2Å were observed in the protein, PD and SF, respectively in comparison to the WT-closed state (Supplementary Fig.4). Additionally, a stable pattern was observed in the C_α_RMSD values of protein (<7Å), PD (<4.2Å) and SF (<4.2Å) in the G300D-closed structure during the MD simulation (Supplementary Fig.4). Similar to the WT-open state, a highly stable pattern of C_α_RMSD values below 5Å is demonstrated by the protein, SF and PD of the R67Q-open, R218L-open and G300D-open structures (Supplementary Fig. 4). Most evidently in comparison to WT-open, R67Q-open, and R218L-open mutant structures higher fluctuations /deviations are observed in the G300D-open structure (mainly in the PD and SF) during the last 150ns of the MD simulation (Supplementary Fig.4). The overall RMSD plot of protein stability in open conformation state is shown in Fig. 6a.

The stability of the systems under investigation was further assessed using the Root-Mean-Square-Fluctuations (RMSF), which is used to assess the mobility/stability of proteins after MD simulations. The RSMF identifies which amino acids within a protein play a significant role in molecular motion by indicating their structural contributions. It quantifies the displacement of a specific atom or group of atoms with respect to the reference structure, and the average is calculated across the total number of atoms^20^. The RMSF was monitored for all the systems with the highest fluctuations observed in the highly plastic regions of the Kir2.1 ion channel, including the extracellular loop regions (residues 25-50, 110-125) and the C-terminal regions (Fig. 6d-k). Overall, TMD1 (residues 75-110), TMD2 (residues 150-182), selectivity filter (residues 141-147) and the G-loop (residues 300-306) regions displayed lower fluctuations (RMSF>3Å). This indicates that these regions exhibit stable backbone conformations in the Kir2.1 channel. RMSF values between 3 to 12Å were observed in the external loop, extracellular loop, and C-terminal region of the WT-closed state, while in other regions lower fluctuations within 1 to 5Å were witnessed for each chain (Fig. 6e). Similarly, higher fluctuations were observed in the external loop, extracellular loop, and C-terminal region of the mutants including R67Q-closed, R218L-closed and G300D-closed (Fig. 6g, i, k). Additionally, in all mutant closed-state structures, high RMSF values up to 6Å were observed for the residues (200 to 250) constituting the solvent-exposed loop (Fig. 6g, i, k).

Higher flexibility (RMSF 1-7Å) was shown by the residues forming the external loop, extracellular loop, solvent exposed loop and C-terminal region in the WT-open state of Kir2.1 (Fig. 6d). Lower backbone flexibility (RMSF>3 Å) was depicted by other regions of the open state Kir2.1 (Fig. 6d). In comparison to WT-open state, very high fluctuations with RMSF values up to 9Å were observed in the highly plastic regions (i.e., external loop, extracellular loop, solvent exposed loop and C-terminal) of the channel when mutations were introduced, including R67Q-open, R218L-open, and G300D-open. However, other regions of R67Q-open, R218L-open, and G300D-open states show less flexibility as indicated by the RMSF values less than 3Å (Fig. 6f, h, j). Fig. 6c depicts the movement of potassium ion from open conformation state of the channel and the segments S0-S4 are the compartments of selectivity filter shown in Fig. 2b conforming the open conducting state of the channel. In total, 5800 frames were extracted from individual simulations. To further investigate the dynamic and dominant motion or conformational changes from trajectories, we performed PCA and NM analysis, which help identifying and focus on the dominant motions by dimensionality reduction.

### Principle Component Analysis and NM analysis

Principle Component Analysis (PCA)^21^ was performed to analyze the structural variance during MD simulation. While different conformational states along the trajectories were studied with representative structures of highly populated clusters (Fig.7), the coordinate covariance matrix from the trajectories and PCs were used to identify and visualize the dominant motion and explain the most significant structural changes and dynamics. The clusters represent certain conformations or modes of motion undergone by structures over time. While a single PC explains a specific transition of the system, all the PCs collectively explain the complete range of motions and conformational changes observed during the simulation. This comprehensive analysis allows for a more thorough understanding of the protein’s dynamic behavior across the entire trajectory.

**Fig. 7:**
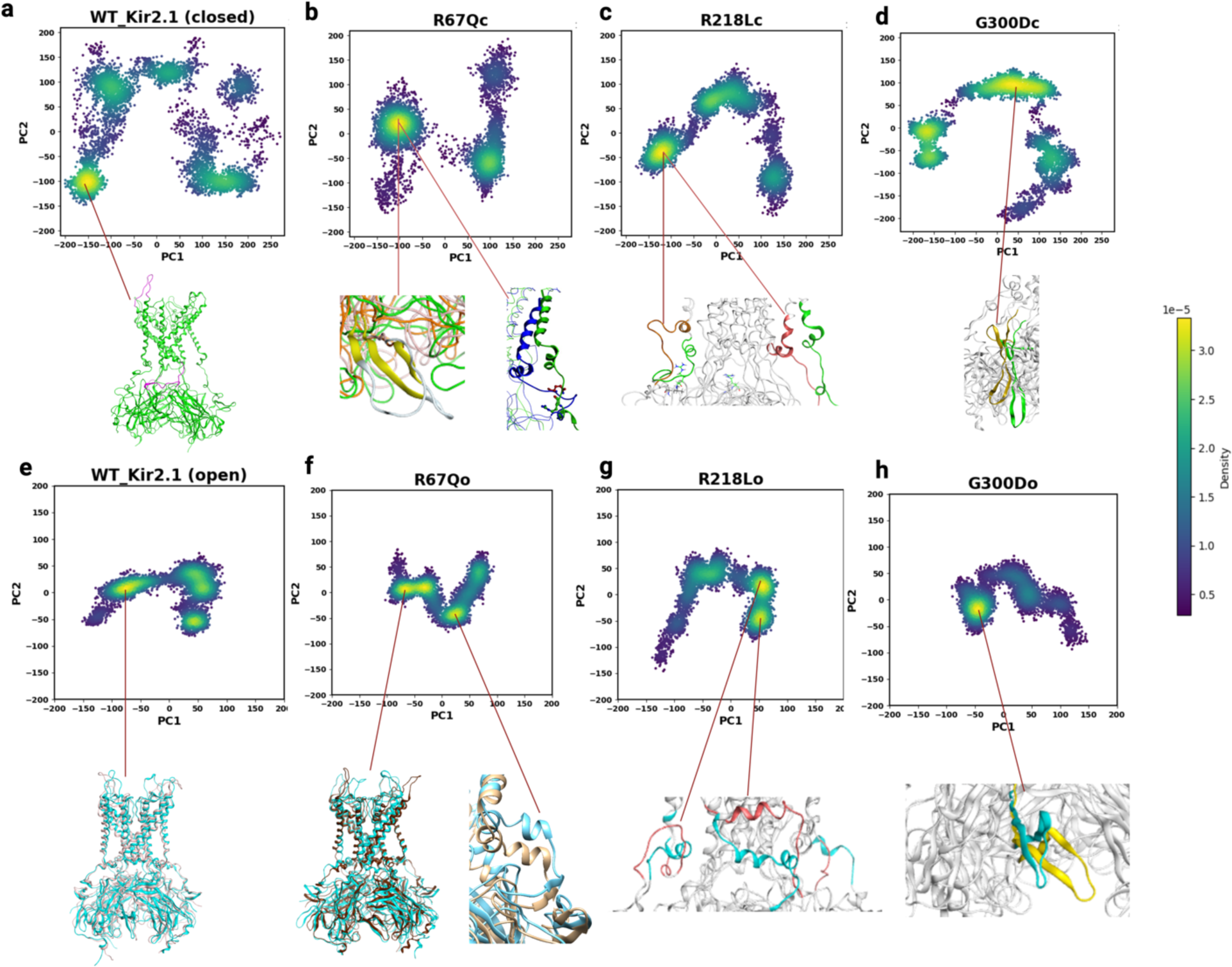
PCA analysis of MD trajectories of Kir2.1-WT and mutants in open and closed conformation the inset underneath each PCA plot is showing the representative frame from local minima and explaining the change with respect to refence structure. **a)** PCA plot explains the projection of molecular dynamics trajectories onto first two principal components (PC1 and PC2). Each point represents a snapshot from the trajectory, and clusters indicate regions of the conformational space where the protein spends significant time, corresponding to energy minima. The color density indicates the population density of the snapshots, with yellow representing higher density and purple representing lower density. for Kir2.1-WT in closed state **b)** PCA plot of R67Q in closed state **c)** PCA plot of R218L in closed state **d)** PCA plot of G300D in closed conformation state. **e)** PCA plot of R67Q in open state **f)** PCA plot of R218L in open state **g)** PCA plot of G300D in closed conformation state. The structure underneath each plot is showing the represented frame structure overlay on refence frame in open conformation state.

In the closed state, WT Kir2.1 exhibited multiple distinct clusters, indicating a range of stable conformations (Fig. 7a). Regions of heightened flexibility were observed in the N-terminal (residues 50-70), extracellular loop (residues 110-130), and slide helix (residues 70-80), consistent with RMSF plots. The slide helix, crucial for the gating mechanism, connects the transmembrane and cytoplasmic domains. The R67Q mutant displayed fewer, more dispersed clusters compared to WT, suggesting altered conformational stability. Structural changes were particularly evident in the slide helix region (Fig.7b). R218L showed a more continuous distribution of states, implying increased flexibility. Notably, this mutation induced alterations in the slide helix position and formation (Fig. 7c). The G300D mutant exhibited a unique pattern with two major clusters, indicating significant changes in conformational behavior. G300D affected both the slide helix position and cytoplasmic loop stability, as shown in the inset (Fig. 7d) and RMSD plot (Fig. 6j).

In the open conformation, WT Kir2.1 demonstrated a more compact distribution compared to its closed state, with flexibility primarily in the extracellular loop region, N and C-termini (Fig. 7e). The R67Q mutant exhibited distinct conformational changes compared to the WT. The most pronounced fluctuations were observed in the extracellular loop region, with notable alterations also seen in the N-terminal portion of the cytoplasmic domain. While the slide helix showed a slight bend, its overall structural stability remained largely intact. Interestingly, the 2D structural features, although mobile, maintained their fundamental formation throughout the simulation (Fig. 7f). This preservation of the secondary structure, despite increased flexibility, suggests a subtle modulation of channel dynamics rather than a dramatic structural reorganization. R218L displayed significant changes in the extracellular and C-terminal cytoplasmic loop regions (residues 200-220), with the slide helix transitioning from a helical to a loop structure. This structural alteration could impact interactions with regulatory lipids such as PIP2 (Fig. 7e). The G300D mutation induced major fluctuations in the cytoplasmic loop region, likely due to the introduction of a negatively charged residue in place of a hydrophobic one. This change causes more deviation of the structure from its position, and the single change at the G-loop starting position subsequently affects the large conformational difference compared to the WT. This PCA analysis provides crucial molecular-level insights into how these ATS-associated mutations alter the structural dynamics of Kir2.1, contributing to our understanding of the mechanistic basis of channelopathy in Andersen-Tawil Syndrome and other Kir2.1-LOF-linked conditions. For further analysis of the principal motions and overall dynamic behavior during the simulation period, NM analysis was performed.

NM analysis, derived from PCA, revealed distinct conformational dynamics in Kir2.1 WT and its mutants (R67Q, R218L, and G300D) in both closed and open states (Fig. 8). In the closed state, Kir2.1-WT exhibited significant flexibility in the N-terminal, extracellular loop, and C-terminal regions, with notable movement of the slide helix crucial for gating (Fig. 8a <u>Supplementary movie Kir2.1-WT_closed</u>). Conversely, the open state demonstrated increased overall stability, with fluctuations primarily localized to extracellular regions (Fig. 8e <u>Supplementary movie Kir2.1-WT_open</u>). These findings corroborate our RMSD and RMSF results, also PCA indicated multiple stable conformations in the closed state and a more compact distribution in the open state.

**Fig. 8.**
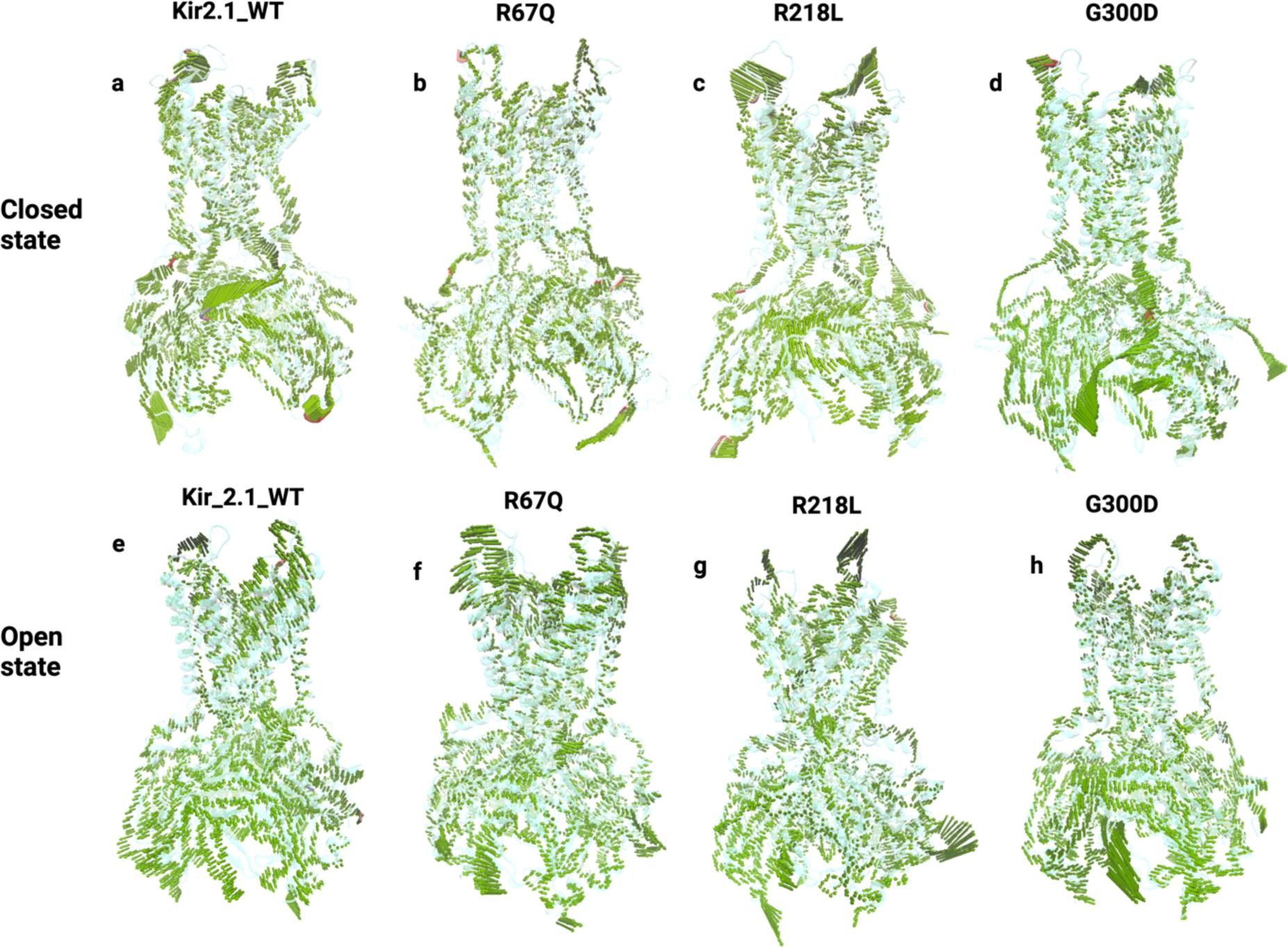
PCA based NM analysis of Kir2.1-WT and ATS-mutations (R67Q, R218L, G300D) in open and closed State. The porcupine plots illustrate the likely modes of large-scale collective motion, direction, and magnitude of atomic displacements along the principal normal modes derived from PCA for a) Kir2.1-WT closed and b) R67Q closed, c) R218L closed, and d) G300D in the closed state. The second row shows e) Kir2.1-WT open state, f) R67Q open, g) R218L open, and h) G300D) in open conformations state. Each green arrow represents the direction and relative magnitude of motion for Ca atoms of each residue.

The R67Q mutation induced subtle changes in the slide helix conformation while maintaining overall structural integrity in the closed state (Fig. 8b <u>Supplementary movie R67Q_closed</u>). Surprisingly, the open state exhibited increased mobility in extracellular and cytoplasmic loops, with tether helix fluctuations potentially compromising channel function (Fig. 8f <u>Supplementary movie R67Q_open</u>). These findings are aligned with RMSD plot and PCA-derived altered conformational stability. Similarly, R218L demonstrated abnormal fluctuations in the extracellular domain and slide helix in the closed state (Fig. 8c <u>Supplementary movie R218L_closed</u>). The open state revealed increased cytoplasmic loop mobility and disrupted tether helix formation, significantly altering the orientation of gating residues (Fig. 8g <u>Supplementary movie R218L_open</u>). This enhanced flexibility was also captured in our PCA, showing a continuous distribution of states. The G300D mutation, involving a charge alteration in the critical G-loop region, induced pronounced changes in atomic displacement patterns in both conformational states (Fig. 8d <u>Supplementary movie G300D_closed</u>, Fig. 8h <u>Supplementary movie G300D_open</u>). Abnormal cytoplasmic loop movements affected G-loop stability and lower gating residues, consistent with PCA-observed distinct conformational landscapes.

## Discussion

In this study, we performed atomic-level investigation of *KCNJ2* clinical mutations R67Q, R218L, and G300D compared to the WT structure. To elucidate the atomic-level mechanisms of *KCNJ2* mutations, we developed the first full-length Kir2.1 model, addressing the limitations of previous truncated structures lacking ∼40 N-terminal and 60-65 C-terminal residues^22^ ^23^ ^24^, or use of the cytoplasmic domain alone^10^. Our approach enabled the inclusion of complete channel architecture, encompassing regions critical for lipid interactions, gating mechanisms, and regulatory regions. The complementary models used allowed for unique detailed features: Site-directed mutagenesis showed local interaction profiles upon specific mutation; molecular dynamics (MD) simulations were utilized to assess overall structural perturbations and dynamic conformational changes; principal component analysis (PCA)^25^ assessed the collective atomic motion of the mutant compared to WT; PCA allowed obtaining large-scale collective motion of atoms that frequently correlates the biological function and biophysical properties^26^, Free energy landscape (FEL)^27^ plots helped identify local minima, with representative frames revealing stable conformational changes; and porcupine plots, constructed using normal mode analysis, illustrated the amplitude and direction of movement in specific structural elements, such as loops and helices. Together, open and closed conformation state in-depth investigations have determined how single residue changes induce large conformation changes to the overall Kir2.1 channel structure, disturbing normal conductance, thus leading to loss of normal function.

To further elucidate the gating mechanism, we compared pore radius profiles of the closed and open state models. This analysis revealed distinct differences in pore dimensions, particularly at the helix bundle crossing (HBC) and G-loop regions. The closed state exhibited a narrower pore, consistent with the role of these regions in restricting ion flow. Intriguingly, we observed a slight difference in pore dimensions at the selectivity filter region between the two states, suggesting subtle conformational changes during gating. The cryo-EM structure of the closed state identified I176, M180, and A306 as key gating residues^6^ (Fig. 2b). The open state model, although based on Kir2.2, indicated that G177 and M180 are also involved in gating. Despite the template difference, the importance of these HBC residues is evident in Kir2.1, emphasizing their role in channel dynamics. In other studies, *KCNJ2* mutation G177E resulted in a forced-open conformation with pH dependence^28^. Interestingly, the extracellular disulfide bond C122 is important for stabilizing the open state of the channel, and the C122Y mutation can alter the hydrogen bond network with PIP2, leading to arrhythmogenesis in ATS^24^. Additionally, G177 in Kir2.1, equivalent to G178 in Kir2.2, has been implicated in subunit gating^28^. These reports also confirm the role of G177 and M307 residues in channel gating. Similarly, the M307V mutation led to a loss-of-function phenotype and increased channel sensitivity to PIP2 depletion^29^. Our findings advance our understanding of the residues involved in Kir2.1 channel gating.

Site-directed mutation analysis identified atomic-level structural interactions that were lacking and new abnormal interactions in the *KCNJ2* mutations. Here, we examined ATS-associated mutations R67Q, R218L, and G300D in both open and closed states, demonstrating significant alterations in local hydrogen bonding networks. Notably, the R67Q mutation in the closed state led to the loss of key interactions and the formation of novel bonds, indicating a structural displacement. In the open state, R67Q disrupted an intricate hydrogen bonding pattern crucial for α-helix stability. The R218L mutation similarly resulted in the loss of dense hydrogen bonding networks in both conformational states, potentially affecting channel gating mechanisms. The G300D mutation, while maintaining some conserved interactions, introduced new hydrogen bonds, particularly with T309, suggesting altered G-loop dynamics. MD analysis further corroborated these findings, revealing dynamic changes in protein structure and interaction patterns over time. This computational approach provided additional insights into the stability of the newly formed interactions and their potential impact on channel function. These findings highlight the complex interplay between single amino acid substitutions and channel architecture, extending beyond local effects to potentially influence global protein stability and function. Similar approaches have been applied elsewhere^30^ and revealed insights into the structural perturbations. Our data expands the observed changes in interaction profiles thus, providing a molecular basis for understanding the functional defects in Kir2.1 mutants and offers new insights into the structural underpinnings of channelopathies.

The MD simulation revealed that the conformational changes induced by mutants influenced loss of overall stability of the channel in the closed state and led to the loss of native Kir2.1 conformation. Previously, simulation studies investigated slide helix conformations during HBC opening^31^. Interestingly, we found that the open conformation state mutant models were comparatively stable. A recent MD simulation report showed that the disruption of the C122/C154 disulfate bond at the extracellular loop region causes structure instability of the selectivity filter and impairs the K^+^ flux^32^. It is postulated that this disrupts the regulatory lipids binding to the channel^24^. Therefore, we specifically mentioned this bond while preparing the lipid-by-layer membrane in the Charmm-gui^33^ for every simulation system. The porcupine plots expand on the explanation for this based on PCA-based NM analysis for R67Q and R218L, in which abnormal fluctuation in the extracellular region, as well as alterations of the side helix stability, cause energetic instability and conduction malfunctioning. By extracting prominent conformations from molecular dynamics trajectories revealed that Kir2.1-WT exhibits movements in the N-terminal, extracellular loop, and slide helix regions. These fluctuations subsequently impact the channel gating and that ATS-associated mutations induce distinct conformational aberrations. The R67Q mutation causes abnormal fluctuations in the slide helix and extracellular loop, potentially disrupting gating transitions. R218L critically compromises the tether helix structure, which leads to impaired pore-cytoplasmic domain coupling. G300D perturbs the G-loop and slide helix conformations, leading to atypical movements in solvent-exposed regions.

The NM analysis was recently used to describe different conformational states in cryoEM structure in the absence of PIP2^6^. However, our study is the first to apply PCA-based NM analysis to *KCNJ2* mutations. Interestingly, conformational changes identified by PCA and NM analysis yielded results consistent with RMSF data. This concordance provides a robust characterization of the structural and dynamic alterations induced by these mutations. This atomic-level analysis not only enhances our understanding of Kir2.1 channel dynamics but also provides a foundation for structure-guided therapeutic strategies targeting specific molecular defects in ATS and *KCNJ2-* related channelopathies.

Our comprehensive structural and functional analysis of *KCNJ2* clinical mutations provides critical insights for developing targeted therapeutic strategies for ATS and related channelopathies. By elucidating specific structural perturbations associated with each mutation, particularly in the slide helix, tether helix, and G-loop regions, our study offers a detailed roadmap for structure-guided drug design. Compounds that can reestablish critical structural connections may rescue channel function, paving the way for mutation-specific, personalized treatments.

### Limitations and Future Directions

Our study of *KCNJ2* mutations in ATS provides valuable insights but we recognize there are some limitations inherent to the nature of this work. Structural analyses were constrained by the available closed-conformation cryo-EM structure of Kir2.1, necessitating the use of a homology model for the open state. Molecular dynamics simulations, while informative, were limited in duration and excluded regulatory molecules like PIP2. Principal component analysis, though valuable for identifying dominant motions, presented challenges in scale-variance and interpretability. In future work, we aim to address these limitations, including developing high-quality experimental models of Kir2.1 in various conformational states, investigating PIP2-channel interactions, and extending simulation times. Additionally, incorporating physiologically relevant heteromeric channel assemblies and validating computational findings through advanced experimental techniques are planned. Further advancements in structural biology and computational methods, particularly in the development of a high-resolution open state Kir2.1, will minimize the limitations of the current model system.

## Material and Methods

### Experimental part

#### Expression Constructs

All missense mutations (R67Q, R218L, G300D) were made using the QuikChange II XL kit using primers designed by the Agilent Primer Design Program (Agilent). The template for mutagenesis was pcDNA3 MYC-tagged WT Kir2.1 previously described^34^. Restriction digest analysis was used to test the integrity of all mutations and mutations were verified by full codon sequencing by the University of Wisconsin Biotechnology Center. Stable cell lines were created by transfecting each construct into HEK 293 cells using lipofectamine 2000 (Invitrogen) and selecting under geneticin. Selected cells were then diluted to isolate individual colonies and screened for expression by western blot and whole-cell patch clamp.

#### Western blotting

HEK 293 cells stably expressing MYC-Kir2.1 (WT, R67Q, R218L, G300D) were detected by SDS-PAGE analysis. Cells were lysed in lysis buffer (50 mM Tris-HCl pH 7.4, 150 mM NaCl, 5mM EDTA, 1% NP40, and Halt Protease Inhibitor Cocktail (Thermo Scientific)) at 4 **°**C. Insoluble material was spun down at 15,000 × *g* for 10 min and supernatants were mixed with an equal amount of Laemmli sample buffer containing urea, separated by SDS/7% PAGE and transferred to PVDF membranes. Membranes were first blocked in blocking buffer (PBS pH 7.5, 0.05% Tween-20 and 5% dry milk) for 20 min then incubated with 1:2000 rabbit anti-Kir2.1 antibody (Santa Cruz Biotechnology) in blocking buffer overnight at 4 **°**C. Membranes were then washed in PBST and incubated in 1:15,000 anti-rabbit HRP secondary antibody (Thermo Scientific) for 1hr before washing again in PBST and detection using Immobilon Western HRP Substrate (Millipore).

#### Whole Cell Patch Clamp recordings Molecular Biology/Cell Culture

Stable Kir2.1-WT and homo-tetrameric mutant cell lines including (R67Q, R218L, and G300D) were cultured in DMEM (Invitrogen) with 10% fetal bovine serum (FBS) and maintained at 37°C in 5% CO2. Cells were prepared 24 h prior to whole cell recordings.

#### Electrophysiology

Whole cell patch clamp experiments were performed at room temperature. Borosilicate glass capillary patch electrodes with resistance 2-4 MΩ when filled (Model P-97, Shutter Instrument, Novata CA). The pipettes were filled with internal solution containing (in μM) K-gluconate 150, EGTA 5, MgATP 5, and HEPES 10 and pH was adjusted to 7.2 with KOH. External bath solution contained (in μM) NaCl 148, KCL 5.4, CaCl2 1.8, MgCl2 1, HEPES 15, NaHPO4 0.4, D-glucose and pH adjusted to 7.4 with NaOH. The whole cell patch clamp mode was applied to the cells in the voltage clamp configuration using Axopatch 200B amplifier (Molecular Devices, Sunnyvale, CA). Current signals were digitized at 10 kHz, filtered at 2 kHz and stored on IBM-CompatiblePC interfaced with a Digidata 1440 analogue to digital converter (Molecular Devices,Sunnyvale, CA). Initial series resistance values were 2-5 MΩ, which was compensated by at least 80% in all experiments. Cell capacitance was performed using acquisition software pClamp 10.2 (Molecular Devices, Sunnyvale, CA). *I_K1_* was recorded starting from a holding potential of −50 mV, voltages were ramped from −120 to +50 mV with a sequential 10 mV steps increase in 100 ms steps. Barium chloride 0.5 mM in bath solution was used to perfuse the cells for a minimum of 2 minutes, and the I-V protocol was repeated. Data analysis was performed using Origin 2020 software (OriginLab, Corporation). Data are expressed as mean ± SE unless otherwise specified. Statistical analysis was performed using one-way ANOVA with Dunnett’s correction for multiple comparisons tests. P values less than 0.005 were considered significant.

### Computational Studies

#### Structural Modeling

The closed-state cryo_EM structure of Kir2.1 was obtained from RCSB^35^ (PDBid: 7zdz). However, the open conformation state of the Kir2.1 was developed using a homology modeling approach. The 427 amino acid long sequence was retrieved from Uniport^36^ (ID: p63253) (https://www.uniprot.org/uniprotkb/P63252/entry). To identify the optimal template an iterative BLASTp search was performed against the Protein Data Bank (PDB) database ^37^. The crystal structure 6M84 with a resolution of 2.81 Å, representing the force open mutant Kir2.2 in complex with PIP2 was selected as the template for homology modeling^38^. Sequence and structure alignment of the target and template was performed using the Align program of MOE software package v2022.02 ^17^ with default settings. For comparative modeling, the tools SWISS-Model ^39^, I-TASSER^40^, MODELLER v10.0^41^ software, and the application implemented in MOE^17^ were used to develop 100 independent models from each software. In MOE following the selection of all loops, the side chain modeling was performed using an extensive rotamer library generated by LowModeMD wherein, systematic clustering of conformations was performed ^42,43^. The models were refined and scored using the using the Coulomb and Generalized Born/ Volume Integral (GB/VI) implicit scoring function. Protonate 3D^44^ was applied prior to model refinement and the Amber10:EHT was used. The top five models were shortlisted based on best GB/VI and modeler scoring. The final model was selected based on model validation and quality assessment using the ERRAT ^45^ and PROCHECK^46^. The full-length open-state homology model for Kir2.1 with the best parameters was selected. Prior to further analysis, the Kir2.1-WT structures and mutant structures (open and closed) were prepared using the energy minimization application in MOE v2022.02^15^ up to 0.1 gradients using the Amber10:EHT forcefield for further modeling studies. The selected mutations were then individually implemented in a homotetramer fashion in the open and closed conformation states, followed by energy minimization to develop the mutant models. The hole pore radius software^16^ was used to generate the central pore in order to validate the channel’s current state and construction/gating residues.

#### Site-directed mutagenesis

The Protein Design module of MOE v2022.02 ^17^ was used for mutation analysis. The inputs for the analysis were prepared using the Residue Scan option whereby a single mutation was inserted at the selected location position in one structure at a time, i.e., GLN at position 67 in all four chains, considering both the open and closed states of Kir2.1. All three open and closed state mutant structures of Kir2.1 were obtained which were further analyzed to elucidate the changes in interactions and stability values compared to the WT-open and WT-closed structures of Kir2.1.

#### Mutational Analysis

After selecting mutations, the analysis was performed on both the open and closed structures of the Kir2.1 ion channel. The Protein Design module of MOE v2022.02 was used for mutation analysis. The inputs for the analysis were prepared using the Residue Scan option, whereby a single mutation was inserted at the appropriate position in all chains of the structure at a time, considering both the open and closed states of Kir2.1. It generates a database with all possible interaction changes on all four chains. These outputs were analyzed to elucidate the changes in interactions and stability values compared to the WT-open and WT-closed structures of Kir2.1. Therefore, the interaction profiles, stability (dStability) and affinity (dAffinity) scores for specific amino acid residues were used to assess relative binding energy changes observed upon the transformation of residues.

#### Molecular Dynamics Simulations

Firstly, open and closed conformation state structures were prepared by adding K^+^ ions and TIP3 water molecules in the selectivity filter and pore region, respectively, to maintain the physiological conditions and stabilization of the selectivity filter. The prepared systems were embedded into the pre-equilibrated 1-palmitoyl-2-oleoyl-sn-glycero-3-phosphocholine (POPC) lipid bilayer membrane using CHARMM-GUI interface ^47^ with 178 lipids on the upper and lower leaflet. Several parameters were introduced at all four chains, including mutations at specific sites and disulfide bonds at residue positions 122 and 154. RUN PPM 2.0 was selected for appropriate orientation and positioning options for all the systems. The simulation cell’s average area and box dimensions were determined to be 12157.40 Å², X = 110.26, Y = 110.26, and Z = 215.38, respectively. The systems were solvated by 0.15 M KCl aqueous solution. In the next step, all formerly built components were assembled, and the systems were equilibrated. A prolonged equilibration protocol for 80ns involving a gradual reduction of restraints was executed (for more information see supplementary Fig.3) on the high-performance computer (HPC) Expanse using Amber22^19^. The Position restraints were imposed on either the WT or mutated complexes or their individual components to maintain system stability during each step in the equilibration molecular dynamics (MD) runs. Additionally, weak restraints on the backbone atoms of the Kir2.1 selectivity filter (SF) were consistently upheld throughout the production runs of 500ns. The production runs spanning a duration of up to 1.5μs, i.e. 580 ns for each replicate, were used for all systems. However, The trajectories were analyzed using in-house Python scripts and the CPPTRAJ module of Amber22 ^19,48^. The stability of each system was assessed by evaluating the C_α_RMSD, C_α_RMSF, Rg, and hydrogen bond analysis with respect to the starting structures.

#### Principal Component Analysis (PCA)

To reduce the dimensionality, we employed Principal Components Analysis (PCA) on trajectory data using the CPPTRAJ program module to explore the conformational dynamics of both wild-type (WT) and mutations. This involves translating the molecule to its average geometrical center and superimposing it onto a reference structure using least squares fitting. Subsequently, we constructed a covariance matrix in Cartesian coordinate space, with dimensions of 3N × 3N, where N represents the number of atoms in the molecule. Diagonalizing this matrix yields a set of eigenvectors, each describing a specific motion component and its corresponding eigenvalue, indicating its energetic contribution. Outputs, including eigenvectors and a trajectory file containing principal modes, were generated for further analysis. A Python script was used to plot and visualize the first two principal components. The resulting graphs showed different clusters representing energy minima identified during simulations. The representative frame from each cluster was identified and turned into PDB files using VMD. Each simulation with WT and mutant was then visually analyzed, and clusters were compared in open and closed state. The mutant clusters were then compared with the WT in both states to inspect global changes in the structure conformation.

#### Normal Mode Analysis

Normal Mode Analysis (NMA) was performed in Cartesian coordinate space with the same parameter set used for the MD. Cartesian coordinates are mass-weighted in the same way as in the principal component analysis. The potential energy was minimized, and the Hessian matrix was then calculated and diagonalized to obtain the eigenvalues and the eigenvector matrix W. The eigenvalues, which are the angular frequencies squared (ω_i_^2^), define the normal modes’ oscillation frequencies. The normal modes themselves are given by the columns of W, which are normalized to the identity matrix. The mean-square fluctuation of the ith atomic coordinate is given by

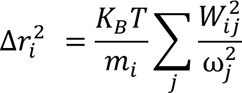

where:

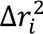 is the mean-square fluctuation of the ith atomic coordinate

- *K_B_* is the Boltzmann constant
- T is the temperature
- *m_i_* is the mass of the ith atom
- 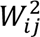 is the (i,j)-th element of the eigenvector matrix W
- ω*_j_* is the angular frequency of the j-th normal mode.

This framework allows for the detailed analysis of the intrinsic motions of the system, providing insights into the dynamic behavior of biomolecules. VMD is used to generate the porcupine plots to assess the direction and magnitude of atomic displacement of each residue.

## Supporting information

R218L_open

R67Q_open

G300D_closed

R218L_Closed

R67Q_closed

Kir2.1_WT_closed

G300D_open

Kir2.1_WT_open

## Notes

### Competing Interest Statement

The authors have declared no competing interest.

## REFERENCES

1. Reilly, L. & Eckhardt, L.L. Cardiac potassium inward rectifier Kir2: Review of structure, regulation, pharmacology, and arrhythmogenesis. Heart Rhythm 18, 1423–1434 (2021).

2. Van Ert, H.A. et al. Flecainide treats a novel KCNJ2 mutation associated with Andersen-Tawil syndrome. HeartRhythm Case Rep 3, 151–154 (2017).

3. Kalscheur, M.M. et al. KCNJ2 mutation causes an adrenergic-dependent rectification abnormality with calcium sensitivity and ventricular arrhythmia. Heart Rhythm 11, 885–94 (2014).

4. Reilly, L. et al. Genetic Loss of IK1 Causes Adrenergic-Induced Phase 3 Early Aderdepolarizations and Polymorphic and Bidirectional Ventricular Tachycardia. Circ Arrhythm Electrophysiol 13, e008638 (2020).

5. Eckhardt, L.L. et al. KCNJ2 mutations in arrhythmia patients referred for LQT testing: a mutation T305A with novel effect on rectification properties. Heart Rhythm 4, 323–9 (2007).

6. Fernandes, C.A. et al. Cryo–electron microscopy unveils unique structural features of the human Kir2. 1 channel. Science Advances 8, eabq8489 (2022).

7. Tao, X., Avalos, J.L., Chen, J. & MacKinnon, R. Crystal structure of the eukaryotic strong inward-rectifier K+ channel Kir2.2 at 3.1 A resolution. Science 326, 1668–74 (2009).

8. Zangerl-Plessl, E.-M. et al. Atomistic basis of opening and conduction in mammalian inward rectifier potassium (Kir2. 2) channels. Journal of General Physiology 152, e201912422 (2019).

9. Donaldson, M. et al. PIP2 binding residues of Kir2. 1 are common targets of mutations causing Andersen syndrome. Neurology 60, 1811-1816 (2003).

10. Pegan, S. et al. Cytoplasmic domain structures of Kir2. 1 and Kir3. 1 show sites for modulating gating and rectification. Nature neuroscience 8, 279-287 (2005).

11. Ma, D., Tang, X.D., Rogers, T.B. & Welling, P.A. An andersen-Tawil syndrome mutation in Kir2. 1 (V302M) alters the G-loop cytoplasmic K+ conduction pathway. Journal of biological chemistry 282, 5781-5789 (2007).

12. Tan, B.-H., Dovat, S., Peterson, B.Z. & Song, C. Reduced PIP2 Binding to KCNJ2 (M307I) Channels is Linked to Type 1 Andersen-Tawil Syndrome. Biophysical Journal 104, 129a (2013).

13. Whorton, M.R. & MacKinnon, R. Crystal structure of the mammalian GIRK2 K+ channel and gating regulation by G proteins, PIP2, and sodium. Cell 147, 199-208 (2011).

14. Whorton, M.R. & MacKinnon, R. X-ray structure of the mammalian GIRK2–βγ G-protein complex. Nature 498, 190–197 (2013).

15. Inc, C.C.G. Molecular operating environment (MOE). (Chemical Computing Group Inc. Montreal, QC, Canada, 2016).

16. Smart, O.S., Neduvelil, J.G., Wang, X., Wallace, B. & Sansom, M.S. HOLE: a program for the analysis of the pore dimensions of ion channel structural models. Journal of molecular graphics 14, 354–360 (1996).

17. ULC, C.C.G. Molecular Operating Environment (MOE)(2022.02). (Chemical Computing Group ULC Montreal, QC, Canada, 2022).

18. Hibino, H. et al. Inwardly rectifying potassium channels: their structure, function, and physiological roles. Physiological reviews 90, 291–366 (2010).

19. Case, D.A. et al. AmberTools. Journal of Chemical Information and Modeling 63, 6183–6191 (2023).

20. Marmnez, L. Automatic identification of mobile and rigid substructures in molecular dynamics simulations and fractional structural fluctuation analysis. PLoS One 10, e0119264 (2015).

21. Greenacre, M., et al. Principal component analysis. Nature Reviews Methods Primers 2, 100 (2022).

22. Yeh, S.-H., Chang, H.-K. & Shieh, R.-C. Electrostatics in the cytoplasmic pore produce intrinsic inward rectification in Kir2. 1 channels. The Journal of general physiology 126, 551-562 (2005).

23. Tao, X., Avalos, J.L., Chen, J. & MacKinnon, R. Crystal structure of the eukaryotic strong inward-rectifier K+ channel Kir2. 2 at 3.1 Å resolution. Science 326, 1668-1674 (2009).

24. Cruz, F.M. et al. Extracellular Kir2. 1C122Y Mutant Upsets Kir2. 1-PIP2 Bonds and Is Arrhythmogenic in Andersen-Tawil Syndrome. Circulation Research 134, e52-e71 (2024).

25. Ichiye, T. & Karplus, M. Collective motions in proteins: a covariance analysis of atomic fluctuations in molecular dynamics and normal mode simulations. Proteins: Structure, Function, and Bioinformatics 11, 205–217 (1991).

26. Amadei, A., Linssen, A.B. & Berendsen, H.J. Essential dynamics of proteins. Proteins: Structure Function, and Bioinformatics 17, 412–425 (1993).

27. Papaleo, E., Meregheq, P., Fantucci, P., Grandori, R. & De Gioia, L. Free-energy landscape, principal component analysis, and structural clustering to identify representative conformations from molecular dynamics simulations: The myoglobin case. Journal of molecular graphics and modelling 27, 889–899 (2009).

28. Maksaev, G. et al. Subunit gating resulting from individual protonation events in Kir2 channels. Nature communications 14, 4538 (2023).

29. Handklo-Jamal, R. et al. Andersen–Tawil syndrome is associated with impaired Pip2 regulation of the potassium channel Kir2. 1. Frontiers in Pharmacology 11, 672 (2020).

30. Kalathiya, U., Padariya, M. & Baginski, M. Structural, functional, and stability change predictions in human telomerase upon specific point mutations. Scientific Reports 9, 8707 (2019).

31. Meng, X., Cui, M. & Logothetis, D.E. Simulations of the Helix Bundle Crossing Gate Opening in Kir Channels. Biophysical Journal 104, 129a (2013).

32. Zuniga, D. et al. Biochemical, biophysical, and structural investigations of two mutants (C154Y and R312H) of the human Kir2. 1 channel involved in the Andersen-Tawil syndrome. bioRxiv, 2024.02. 09.579451 (2024).

33. Lee, J. et al. CHARMM-GUI input generator for NAMD, GROMACS, AMBER, OpenMM, and CHARMM/OpenMM simulations using the CHARMM36 additive force field. Biophysical journal 110, 641a (2016).

34. Brown, K.A. et al. Proteomic analysis of the functional inward rectifier potassium channel (Kir) 2.1 reveals several novel phosphorylation sites. Biochemistry 60, 3292–3301 (2021).

35. Rose, P.W. et al. The RCSB Protein Data Bank: redesigned web site and web services. Nucleic acids research 39, D392–D401 (2010).

36. Bateman, A. et al. UniProt: the universal protein knowledgebase in 2023. Nucleic Acids Research 51(2022).

37. Altschul, S.F., Gish, W., Miller, W., Myers, E.W. & Lipman, D.J. Basic local alignment search tool. Journal of molecular biology 215, 403–410 (1990).

38. Zangerl-Plessl, E.-M. et al. Atomistic basis of opening and conduction in mammalian inward rectifier potassium (Kir2. 2) channels. Journal of General Physiology 152(2020).

39. Waterhouse, A. et al. SWISS-MODEL: homology modelling of protein structures and complexes. Nucleic acids research 46, W296–W303 (2018).

40. Yang, J. et al. The I-TASSER Suite: protein structure and function prediction. Nature methods 12, 7–8 (2015).

41. Webb, B. & Sali, A. Comparative protein structure modeling using MODELLER. Current protocols in bioinformatics 54, 5.6. 1–5.6.37 (2016).

42. Labute, P. LowModeMDD implicit low-mode velocity filtering applied to conformational search of macrocycles and protein loops. Journal of chemical information and modeling 50, 792–800 (2010).

43. Friedrich, N.-O. et al. Benchmarking commercial conformer ensemble generators. Journal of chemical information and modeling 57, 2719–2728 (2017).

44. Labute, P. Protonate 3D: assignment of macromolecular protonation state and geometry. Chemical Computing Group Inc (2007).

45. Colovos, C. & Yeates, T.O. Verification of protein structures: paserns of nonbonded atomic interactions. Protein science 2, 1511–1519 (1993).

46. Laskowski, R.A., MacArthur, M.W., Moss, D.S. & Thornton, J.M. PROCHECK: a program to check the stereochemical quality of protein structures. Journal of applied crystallography 26, 283–291 (1993).

47. Jo, S., Kim, T., Iyer, V.G. & Im, W. CHARMM-GUI: a web-based graphical user interface for CHARMM. Journal of computational chemistry 29, 1859–1865 (2008).

48. Roe, D.R. & Cheatham III, T.E. PTRAJ and CPPTRAJ: software for processing and analysis of molecular dynamics trajectory data. Journal of chemical theory and computation 9, 3084–3095 (2013).

