## Supplementary_information for "Atomic-Level Investigation of *KCNJ2* Mutations Associated with Ventricular Arrhythmic Syndrome Phenotypes"

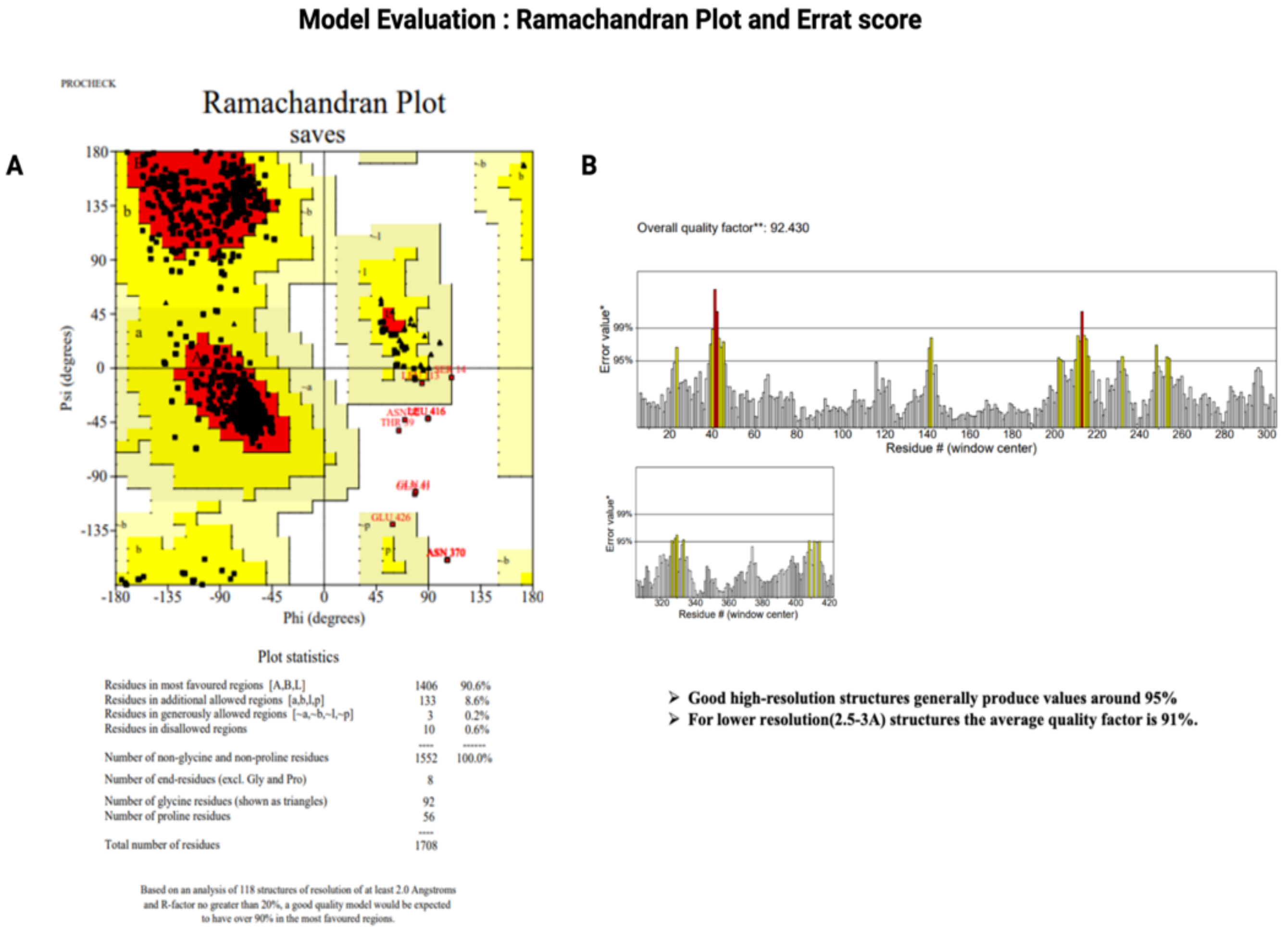


**Supplementary Fig.1: Model Evaluation parameters of the final model**: A) Ramachandran plot B) Errat score plot

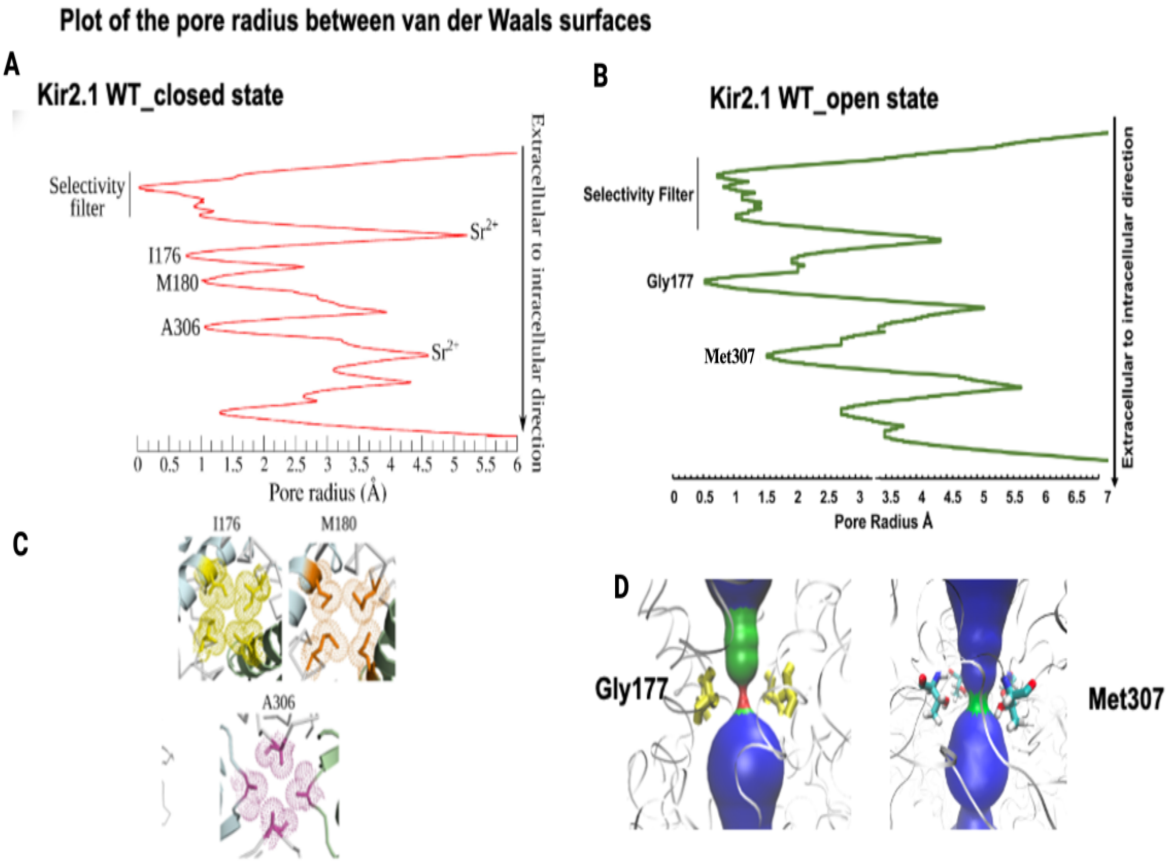


**Supplementary Fig.2: Pore radius plot showing pore area in Å in extracellular to intra cellular direction**: A) closed conformation state pore radius plot and gating residues taken from cryo_EM article of Kir2.1[1], B) the pore radius plot of the open conformation state highlighting the additional Gly177 and Met307 gating residues, C and D captions are missing

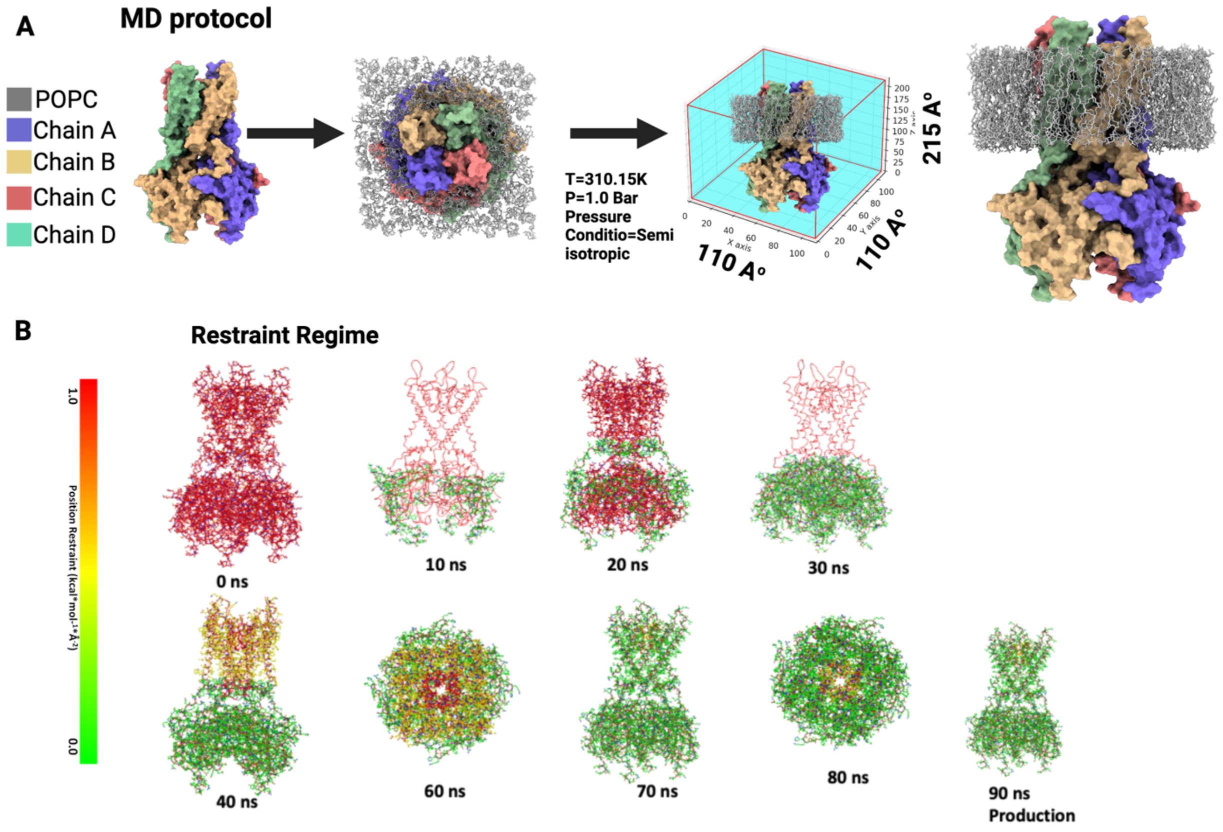


**Supplementary Fig.3: MD protocol and Extended Restraint Regime for MD Simulation: A)** MD conditions and simulation box size also showing the POPC location, **B)** An 80 ns long equilibration protocol using Amber22 on the high-performance computer cluster (HPC) Expanse provides stable MD simulations. The protocol involves varying the location and strength of the restraints applied to the Kir2.1 tetramer at each time point indicated in nanoseconds (ns) and is shaded in light green. This was followed by a mostly unrestrained production run.

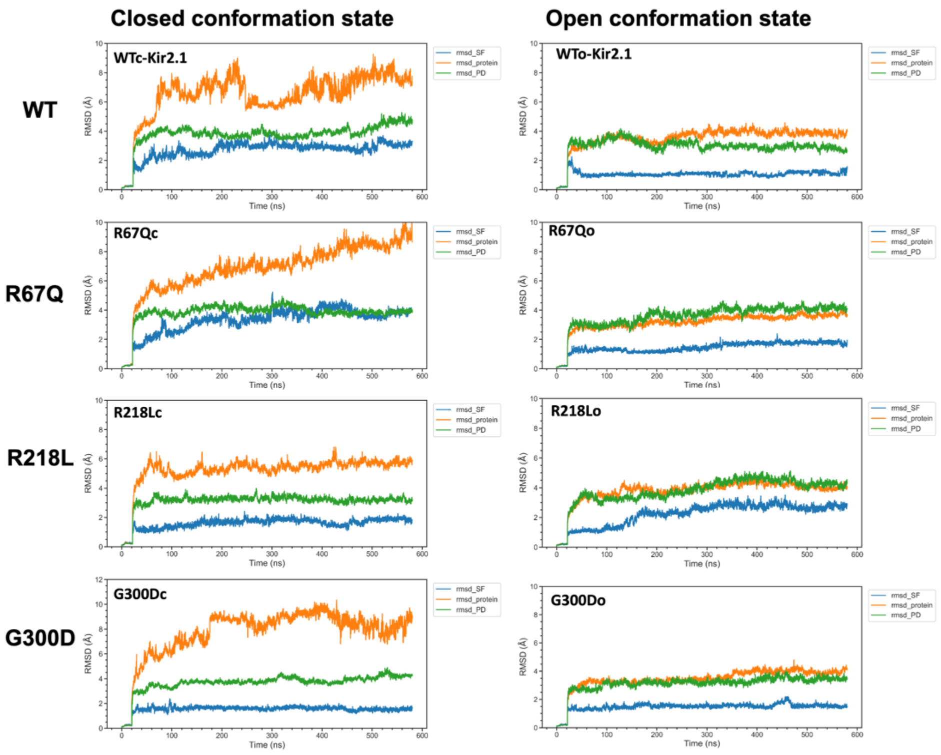


**Supplementary Fig.4:** The CαRMSD values of the WT-closed, WT-open and mutant structures of the Kir2.1 ion channel. The left panel shows the $C_{\alpha}$RMSD plots of the WT-closed, R67Q-closed (R67Qc), R218L-closed (R218Lc) and G300D-closed (G300Dc) mutants while the right panel (top to bottom) shows the $C_{\alpha}$RMSD plots of the WT-open, R67Q-open (R67Qo), R218L-open (R218Lo) and G300D-open (G300Do) structures of Kir2.1. The $C_{\alpha}$RMSD plots for the protein, SF and PD are presented separately in each plot.

**Supplementary Table 1: The detailed interaction pattern of the residues in the WT and mutant open and closed states of the Kir2.1 channel.**

| **CLOSED CONFORMATION OF Kir2.1<br>MUTATION: R67Q (ARG67GLN)** | | |
|---|---|---|
| **STATE** | **Interaction Type and Residues** | **Distance** |
| Kir2.1-WT_CLOSED | Hydrogen bond<br>2:LYS187 (HZ2-O) 1:ALA70 | 2.033 |
| R67Q_CLOSED | Hydrogen bond<br>2:LYS187(HZ2-O)1:ALA70 | 2.032 |
| **Kir2.1-WT_CLOSED** | **Hydrogen bond<br>2:GLY65 (O-HH2) 3:ARG218** | **1.806** |
| | **Hydrogen bond<br>3:ARG218 (HH21-OE1) 3:GLU191** | **2.163** |
| | **Hydrogen bond<br>3:ARG218 (HH21-OE2) 3:GLU191** | **1.920** |
| | **Hydrogen bond<br>3:ARG218 (HE-OE2) 3:GLU191** | **1.759** |
| | **Hydrogen bond<br>2:ARG67(HH12-O)2:THR74** | **2.184** |
| | **Hydrogen bond<br>2:ASP71(OD2-HG1) 2:THR74** | **1.744** |
| | **Hydrogen bond<br>2: ASP71(OD2-H)2:THR74** | **1.787** |
| | **Hydrogen bond<br>2:ASP71(O-HG)2:THR75** | **1.819** |
| **R67Q_CLOSED** | **Hydrogen bond<br>2:GLN67(HE21-OD1) 2:ASP71** | **2.143** |
| | **Hydrogen bond<br>2:GLN (O-H)2:LEU69** | **1.899** |
| | **Hydrogen bond<br>2:THR74(OG1-OD1) 2:ASP71** | **2.635** |
| | **Hydrogen bond<br>2:THR74(H-OD2) 2:ASP71** | **1.949** |
| | **Hydrogen bond<br>2:THR74(HG1-OD1) 2:ASP71** | **1.658** |
| | **Hydrogen bond<br>2: THR75(HG1-O) 2:ASP71** | **1.926** |
| | **Hydrogen bond<br>2:GLY65(O-HH22)3:ARG218** | **1.811** |
| | **Hydrogen bond<br>3:GLU191(OE2-HH21)3:ARG218** | **1.685** |
| | **Hydrogen bond<br>3lGLU191(OE1-HE)3:ARG218** | **1.773** |
| Kir2.1-WT_CLOSED | Hydrogen bond | 2.162 |

| | | |
|---|---|---|
| | 3:ARG67(HE-OD1)3:ASP71 | |
| | Hydrogen bond<br>3:ARG67(HH11-OD1)3:ASP71 | 1.695 |
| | Hydrogen bond<br>3:ASP71(H-OG1)3:THR75 | 1.983 |
| R67Q_CLOSED | Hydrogen bond<br>3:ASP71(H-OG1)3:THR75 | 1.982 |
| | Hydrogen bond<br>3:TYR68(H-OE1)3:GLN67 | 1.851 |
| WT_CLOSED | Hydrogen bond<br>4:ASP71(OD2-CA) 4:TYR68 | 3.403 |
| | H-PI INTERATION<br>4:ARG67(CB-PHRNYLRING)<br>4:TYR68 | |
| R67Q_CLOSED | NO INTERACTIONS<br>OBSERVED | |
| **OPEN CONFORMATION OF Kir2.1<br>MUTATION: R67Q** | | |
| **STATE** | **Interaction Type and Residues** | **Distance** |
| **KIR2.1-WT_OPEN** | **Hydrogen bond<br>1: ARG67(HH22-OE2) 3:GLU191** | **2.027** |
| | **Hydrogen bond<br>1:ARG67(HH12-OE1)3:GLU191** | **1.703** |
| | **Hydrogen bond<br>1:ARG67(HH11-O)3:GLU191** | **1.911** |
| | **Hydrogen bond<br>1:ARG67(CD-O)3:GLU191** | **3.320** |
| | **Hydrogen bond<br>1:ARG67(H-O) GLU63** | **2.643** |
| | **Hydrogen bond<br>3:THR192(HG1-OD1)1:ASP71** | **1.880** |
| | **Hydrogen bond<br>1:GLU63(H-OE1)1:GLU63** | **1.960** |
| | **Hydrogen bond<br>3:LEU217(O-HH)1:TYR68** | **1.723** |
| **R67Q_OPEN** | **Hydrogen bond<br>3:THR192(HG1-OD1)ASP71** | **1.915** |
| | **Hydrogen bond<br>3:LEU217(O-HH)1:TYR68** | **1.773** |
| | **Hydrogen bond<br>1:GLU63(OE1-H)1:GLU63** | **1.958** |
| | **H-PI interaction<br>TYR68(CD2-CB)1:LYS64** | |
| | **1:GLN(O-H)1:ASP71** | **1.899** |
| KIR2.1-WT_OPEN | Hydrogen bond<br>1:TYR68(HH-O)3:LEU217 | 1.733 |
| | Hydrogen bond<br>1:GLU63(OE1-H)1:GLU63 | 1.951 |
| | Hydrogen bond<br>1:ASP71(OD1-HG1)3:THR192 | 1.892 |

| | | |
|---|---|---|
| | Hydrogen bond<br>3:GLU191(O-HH11)1:ARG67 | 1.903 |
| | Hydrogen bond<br>3: GLU191(O-CD)1:ARG67 | 3.345 |
| | Hydrogen bond<br>1: ARG67(HH12-OE1)3:GLU191 | 1.697 |
| | Hydrogen bond<br>1: ARG67(HH22-OE2)3:GLU191 | 2.134 |
| R67Q_OPEN | Hydrogen bond<br>3:THR192(HG1-OD1)1:ASP71 | 1.865 |
| | Hydrogen bond<br>1:GLU63(OE1-H) 1:GLU63 | 1.952 |
| | Hydrogen bond<br>1:VAL61(O-CE)1:LYS64 | 3.267 |
| | Hydrogen bond<br>1:TYR68(HH-O)3:LEU217 | 1.769 |
| | H-PI INTERACTION<br>1:LYS64-1:TYR68 | |
| KIR2.1-WT_OPEN | Hydrogen bond<br>1:GLU63(OE1-H)1:GLU63 | 1.839 |
| | Hydrogen bond<br>1:TYR68(HH-O)1:LEU217 | 1.717 |
| | Hydrogen bond<br>3:THR192(HG1-OD1)1:ASP71 | 1.802 |
| | Hydrogen bond<br>3:GLU191(O-CD)1:ARG67 | 3.402 |
| | Hydrogen bond<br>1:ARG67(HH11-O)3:GLU191 | 1.856 |
| | Hydrogen bond<br>1:ARG67(HH12-OE1)3:GLU191 | 1.697 |
| | Hydrogen bond<br>1:ARG67(HH22-OE1)3:GLU191 | 2.463 |
| | Hydrogen bond<br>1:ARG67(HH22-OE2) 3:GLU191 | 2.143 |
| R67Q_OPEN | Hydrogen bond<br>1:ASP71(OD1-HG1)3:THR192 | 1.779 |
| | Hydrogen bond<br>1:GLU63(OE1-H)1:GLU63 | 1.864 |
| | Hydrogen bond<br>3:LEU217(O-HH)1:TYR68 | 1.758 |
| | H-PI interaction<br>1:LYS64-1:TYR68 | |
| KIR2.1-WT_OPEN | Hydrogen bond<br>1:ARG67(HH12-OE1)3:GLU191 | 1.709 |
| | Hydrogen bond<br>1:ARG67(HH22-OE2)3:GLU191 | 1.941 |
| | Hydrogen bond<br>1:ARG67(HH11-O)3:GLU191 | 1.870 |
| | Hydrogen bond<br>1:ARG67(CD-O)3:GLU191 | 3.376 |

| | | |
|---|---|---|
| | Hydrogen bond 1:ASP71(OD1-HG1)3:THR192 | 1.787 |
| | Hydrogen bond 1:GLU63(OE1-H) 1:GLU63 | 1.843 |
| | Hydrogen bond 3:LEU217(O-HH)1:TYR68 | 1.712 |
| R67Q_OPEN | Hydrogen bond 1:GLU63(OE1-H)1:GLU63 | 2.018 |
| | Hydrogen bond 1:VAL61(O-CE)1:LYS64 | 3.334 |
| | Hydrogen bond 3:LEU217(O-HH)1:TYR68 | 1.749 |
| | H-PI interaction 1:LYS64-1:TYR68 | |
| | Hydrogen bond 3:THR192(HG1-OD1)1:ASP71 | 1.781 |
| **CLOSED CONFORMATION OF Kir2.1 MUTATION: R218L** | | |
| **STATE** | **Interaction Type and Residues** | **Distance** |
| **WT_CLOSED** | **Hydrogen bond 1:ARG218 (HH21-OG)1:SER220** | **2.039** |
| | **Hydrogen bond 1:ARG218 (HE-OG)1:SER220** | **2.462** |
| | **Hydrogen bond 1:ARG218 (HE-OG1)1:THR192** | **2.304** |
| | **Hydrogen bond 1:ARG218 (H-OD1)1:ASN216** | **2.366** |
| | **Hydrogen bond 1:SER220(HG-O)1:SER220** | **2.159** |
| | **Hydrogen bond 1:THR192(HG1-OD1)1:ASN216** | **2.030** |
| | **Hydrogen bond 1:SER220(O-HD21)1:ASN216** | **1.761** |
| | **Hydrogen bond 1:LEU217(O-HZ3)4:LYS64** | **2.225** |
| **R218L_CLOSED** | **Hydrogen bond 1:LEU218(H-OD1)1:ASN216** | **2.298** |
| | **Hydrogen bond 1:SER220(HG-O)1:SER220** | **2.121** |
| | **Hydrogen bond 1:SER220(O-HD21)1:ASN216** | **1.745** |
| | **Hydrogen bond 4:LYS64(O-HE)4:ARG67** | **2.183** |
| WT_CLOSED | Hydrogen bond 1:ARG218(HH21-O)1:LEU217 | 2.264 |
| | Hydrogen bond 1:ARG218(HE-O)1:LEU217 | 2.204 |
| | Hydrogen bond 1:ARG218(H-OD1)1:ASN216 | 2.026 |
| | Hydrogen bond | 1.890 |

| | | |
|---|---|---|
| | 1:ASN216(HD21-O)1:SER220 | |
| 218L_CLOSED | Hydrogen bond<br>1:LEU218(H-OD1)1:ASN216 | 2.029 |
| | Hydrogen bond<br>1:SER220(O-HD21)1:ASN216 | 1.929 |
| WT_CLOSED | Hydrogen bond<br>1:ARG218(HH21-OG)1:SER220 | 1.863 |
| | Hydrogen bond<br>1:ARG218(HH11-OE1)1:GLU191 | 1.810 |
| | Hydrogen bond<br>1:SER220(HG-OG1)1:THR192 | 1.783 |
| | Hydrogen bond<br>1:THR192(HG1-O)1:GLU191 | 1.645 |
| | Hydrogen bond<br>1:THR192(O-HD22)1:ASN216 | 1.870 |
| R218L_CLOSED | Hydrogen bond<br>1:LYS188(HZ1-OE1)1:GLU191 | 1.673 |
| | Hydrogen bond<br>1:GLU191(OE2-H)1:GLU191 | 2.113 |
| | Hydrogen bond<br>1:GLU191(O-HG1)1:THR192 | 1.638 |
| | Hydrogen bond<br>1:THR192(OG1-HG)1:SER220 | 1.871 |
| | Hydrogen bond<br>1:THR192(O-HD22)1:ASN216 | 1.880 |
| WT_CLOSED | Hydrogen bond<br>1:ARG218(HE-OG1)1:THR192 | 1.892 |
| | Hydrogen bond<br>1:ARG218(O-HD22)1:ASN216 | 2.305 |
| | Hydrogen bond<br>1:LEU217(O-HZ3)4:LYS64 | 2.019 |
| | Hydrogen bond<br>1:THR192(HG1-O)1:THR192 | 2.145 |
| R218L_CLOSED | Hydrogen bond<br>4:LYS64(HZ3-O)1:LEU217 | 1.849 |
| | Hydrogen bond<br>1:ASN216(HD22-O)1:LEU218 | 1.966 |
| **CLOSED CONFORMATION OF Kir2.1<br>MUTATION: R218L** | | |
| **STATE** | **Interaction Type** | **Distance** |
| **KIR2.1-WT_OPEN** | **Hydrogen bond<br>1:LEU217(O-HH)4:TYR68** | **1.765** |
| | **H-PI INTERACTION<br>1:LYS219-4:PHE58** | |
| | **Hydrogen bond<br>1:ARG189(HE-OG1)1:THR192** | **2.218** |
| | **Hydrogen bond<br>1:ARG218(H-OD1)1:ASN216** | **1.906** |
| | **Hydrogen bond<br>1:ASN216(HD21-O)SER220** | **1.942** |

| | | |
|---|---|---|
| | **Hydrogen bond THR192(HG1-OD1)ASP71** | **1.816** |
| | **Hydrogen bond ARG218(HE-OD1)ASP71** | **1.805** |
| | **Hydrogen bond ARG218(HH11-OD1)ASP71** | **2.070** |
| | **Hydrogen bond ARG218(HH12-OG1)THR75** | **1.824** |
| | **Hydrogen bond THR75(HG1-O)TYR68** | **2.154** |
| | **Hydrogen bond ASP78(OD2-HG)SER220** | **1.665** |
| | **Hydrogen bond THR74(HG1-OD2)ASP71** | **1.648** |
| **R218L_OPEN** | **Hydrogen bond LEU217(O-HH) TYR68** | **1.777** |
| | **H-PI INTERACTION LYS219-PHE58** | |
| | **Hydrogen bond LEU218(H-OD1)ASN216** | **1.918** |
| | **Hydrogen bond ASN216(HD21-O)SER220** | **1.958** |
| | **Hydrogen bond SER220(HG-OD2)ASP78** | **1.664** |
| | **Hydrogen bond ASP71(OD2-HG1)THR74** | **1.621** |
| | **Hydrogen bond ASP71(OD1-HG1)THR192** | **1.698** |
| | **Hydrogen bond ASP71(OG1-HE)ARG189** | **3.532** |
| | **Hydrogen bond ARG189(HE-OG1)THR192** | **2.124** |
| KIR2.1-WT_OPEN | Hydrogen bond 4:THR75(HG1-O)4:TYR68 | 2.156 |
| | Hydrogen bond 4:THR75(OG1-HH12)1:ARG218 | 1.816 |
| | Hydrogen bond 1:ARG218(HH11-OD1)4:ASP71 | 2.108 |
| | Hydrogen bond 1:ARG218(HE-OD1)4:ASP71 | 1.796 |
| | Hydrogen bond 4:ASP71(OD2-HG1)4:THR74 | 1.645 |
| | Hydrogen bond 4:ASP71(OD1-HG1)1:THR192 | 1.819 |
| | Hydrogen bond 1:THR192(OG1-HE)1:ARG189 | 2.206 |
| | Hydrogen bond 4:TYR68(HH-O)1:LEU217 | 1.793 |
| | Hydrogen bond 1:ARG218(H-OD1)1:ASN216 | 1.873 |

| | | |
|---|---|---|
| | Hydrogen bond 1:ASN216(HD21-O)1:SER220 | 1.966 |
| | Hydrogen bond 1:SER220(HG-OD2)4:ASP78 | 1.676 |
| R218L_OPEN | Hydrogen bond 4:ASP71(OD2-HG1)4:THR74 | 1.611 |
| | Hydrogen bond 4:ASP71(OD1-CD)1:ARG189 | 3.527 |
| | Hydrogen bond 4:ASP71(OD1-HG1)1:THR192 | 1.697 |
| | Hydrogen bond 1:THR192(OG1-HE)1:ARG189 | 2.107 |
| | Hydrogen bond 1:ARG189(HH22-O)1:SER220 | 2.460 |
| | Hydrogen bond 1:SER220(O-HD21)1:ASN216 | 1.990 |
| | Hydrogen bond 1:LEU218(H-OD1)1:ASN216 | 1.897 |
| | Hydrogen bond 1:LEU217(O-HH)4:TYR68 | 1.755 |
| | Hydrogen bond 1:SER220(HG-OD2)4:ASP78 | 1.757 |
| KIR2.1-WT_OPEN | Hydrogen bond 4:TYR68(O-HG1)4:THR75 | 2.154 |
| | Hydrogen bond 4:ASP71(OD2-HG1)4:THR74 | 1.675 |
| | Hydrogen bond 4:ASP71(OD1-HG1)1:THR192 | 1.877 |
| | Hydrogen bond 1:THR192(OG1-HE)1:ARG189 | 2.294 |
| | Hydrogen bond 4:ASP78(OD1-HG)1:SER220 | 1.629 |
| | Hydrogen bond 1:SER220(O-HD21)1:ASN216 | 1.974 |
| | Hydrogen bond 1:ASN216(OD1-HH21)1:ARG189 | 2.193 |
| | Hydrogen bond 1:LEU217(O-HH)4:TYR68 | 1.752 |
| | Hydrogen bond 1:ARG218(HH12-OG1)4:THR75 | 1.827 |
| | Hydrogen bond 1:ARG218(HH11-OD1)2:ASP71 | 2.012 |
| | Hydrogen bond 1:ARG218(HE-OD1)4:ASP71 | 1.765 |
| | Hydrogen bond 1:ARG218(H-OD1)1:ASN216 | 1.867 |
| R218L_OPEN | Hydrogen bond 4:THR74(HG1-OD2)4:ASP71 | 1.659 |
| | Hydrogen bond 4:ASP71(O-HG1)1:THR75 | 2.386 |

| | | |
|---|---|---|
| | Hydrogen bond 4:ASP71(OD1-HG1)1:THR192 | 1.750 |
| | Hydrogen bond 1:THR192(OG1-HE)1:ARG189 | 2.138 |
| | Hydrogen bond 4:TYR68(HH-O)1:LEU217 | 1.747 |
| | Hydrogen bond 4:ASP78(OD1-HG)1:SER220 | 1.631 |
| | Hydrogen bond 1:SER220(O-HH22)1:ARG189 | 2.511 |
| | Hydrogen bond 1:SER220(O-HD21)1:ASN216 | 1.966 |
| | Hydrogen bond 1:ASN216(OD1-H)1:LEU218 | 1.888 |
| KIR2.1-WT_OPEN | Hydrogen bond 4:THR75(HG1-O)4:TYR68 | 2.155 |
| | Hydrogen bond 4:THR75(OG1-HH12)1:ARG218 | 1.815 |
| | Hydrogen bond 1:ARG218(HH11-HG1)1:THR192 | 3.357 |
| | Hydrogen bond 1:ARG218(HE-OD1)1:ASP71 | 1.788 |
| | Hydrogen bond 1:ARG218(HE-OD1)1:ASP216 | 1.868 |
| | Hydrogen bond 4:ASP71(OD2-HG1)4:THR74 | 1.671 |
| | Hydrogen bond 1:THR192(OG1-HE)1:ARG189 | 2.256 |
| | Hydrogen bond 4:TYR68(HH-O)1:LEU217 | 1.736 |
| | Hydrogen bond 1:ASN216(HD21-O)1:SER220 | 2.958 |
| | Hydrogen bond 1:SER220(HG-OD2)4:ASP78 | 1.660 |
| | Hydrogen bond 4:ASP71(OD1-HG1)1:THR192 | 1.860 |
| R218L_OPEN | Hydrogen bond 1:THR74(HG1-OD2)4:ASP71 | 1.648 |
| | Hydrogen bond 4:ASP71(OD1-HG1)1:THR192 | 1.746 |
| | Hydrogen bond 1:THR192(OG1-HE)1:ARG189 | 2.171 |
| | Hydrogen bond 4:TYR68(HH-O)1:LEU217 | 1.741 |
| | Hydrogen bond 1:LEU218(H-OD1)1:ASN216 | 1.923 |
| | Hydrogen bond 1:ARG189(HH2-O)1:SER220 | 2.516 |
| | Hydrogen bond 1:SER220(O-HD21)1:ASN216 | 1.979 |

| | | |
|---|---|---|
| | Hydrogen bond<br>1:SER220(HG-OD2)4:ASP78 | 1.649 |
| **OPEN CONFORMATION OF Kir2.1<br>MUTATION: G300D** | | |
| **STATE** | **Interaction Type** | **Distance** |
| **KIR2.1-WT_OPEN** | **Hydrogen bond<br>3:GLU224(OE1-H)1:GLY300** | **1.939** |
| | **Hydrogen bond<br>3:GLU224(H-OE1)1:GLU299** | **1.857** |
| | **Hydrogen bond<br>1:THR308(O-HG1)1:THR208** | **1.807** |
| | **Hydrogen bond<br>1:THR309(H-O)1:GLY300** | **1.910** |
| **G300D_OPEN** | **Hydrogen bond<br>1:THR308(O-HG1)1:THR208** | **1.765** |
| | **Hydrogen bond<br>3:GLU224(H-OE1)1:GLU299** | **1.845** |
| | **Hydrogen bond<br>1:THR309(O-H)1:ASP300** | **2.336** |
| | **Hydrogen bond<br>1:THR309(H-O)1:ASP300** | **2.125** |
| KIR2.1-WT_OPEN | Hydrogen bond<br>4:GLU224(OE1-H):2:GLY300 | 1.982 |
| | Hydrogen bond<br>4:GLU224(H-OE1)2:GLU299 | 1.928 |
| | Hydrogen bond<br>2: GLU224(OE2-HE2)2:HIE226 | 1.821 |
| | Hydrogen bond<br>2:THR309(H-O)2:GLY300 | 1.871 |
| G300D_OPEN | Hydrogen bond<br>4:GLU224(H-OE1)2:GLU299 | 1.915 |
| | Hydrogen bond<br>2: GLU224(OE2-HE2)2:HIE226 | 1.832 |
| | Hydrogen bond<br>1:THR309(O-H)1:ASP300 | 2.92 |
| | Hydrogen bond<br>1:THR309(H-O)1:ASP300 | 1.778 |
| KIR2.1-WT_OPEN | Hydrogen bond<br>3: GLU229(OE1-HE2)2:HIE226 | 1.843 |
| | Hydrogen bond<br>3: GLU229(OE2-H)2:GLU224 | 1.942 |
| | Hydrogen bond<br>2:GLU224(OE1-H)3:ASP300 | 1.912 |
| | Hydrogen bond<br>3:THR309(H-O)3:ASP300 | 1.907 |
| G300D_OPEN | Hydrogen bond<br>3: GLU229(OE1-HE2)2:HIE226 | 1.859 |
| | Hydrogen bond<br>2:GLU224(H-OE2)3:GLU299 | 2.005 |
| | Hydrogen bond | 1.778 |

| | | |
|---|---|---|
| | 3:THR309(H-O)3:ASP300 | |
| | Hydrogen bond<br>3:THR309(O-H)3:ASP300 | 2.36 |
| KIR2.1-WT_OPEN | Hydrogen bond<br>1:GLU224(OE1-HG1)4:THR308 | 2.055 |
| | Hydrogen bond<br>1:GLU224(H-OE1)4:GLU229 | 1.878 |
| | Hydrogen bond<br>4:THR309(O-H)4:GLY300 | 2.020 |
| | Hydrogen bond<br>1:GLU224(H-OE1)4:GLU229 | 1.913 |
| | Hydrogen bond<br>4:THR308(HG1:O)4:ASP300 | 1.673 |
| | Hydrogen bond<br>4:THR309(H-O)4:ASP300 | 2.286 |
| | Hydrogen bond<br>4:THR309(O-H)4:ASP300 | 2.531 |
| **CLOSED CONFORMATION OF Kir2.1<br>MUTATION: G300D** | | |
| **STATE** | **Interaction Type** | **Distance** |
| **WT_CLOSED** | **Hydrogen bond<br>1:GLU224(OE2-) 1:GLU224** | **3.041** |
| | **Hydrogen bond<br>1:VAL302(O-SD)1:MET301** | **3.74** |
| | **Hydrogen bond<br>1:THR309(HG1-O)1:GLY300** | **1.903** |
| | **Hydrogen bond<br>1:THR309(O-H)1:GLY300** | **1.986** |
| | **Hydrogen bond<br>1:THR309(H-O)1:GLY300** | **2.480** |
| | **Hydrogen bond<br>1:THR308(HG1-O)1:THR308** | **1.996** |
| **G300D_CLOSED** | **Hydrogen bond<br>1:GLU224(OE2-) 1:GLU224** | **3.025** |
| | **Hydrogen bond<br>1:VAL302(O-SD)1:MET301** | **3.665** |
| | **Hydrogen bond<br>1:THR309(HG1-OD2)1:ASP300** | **1.623** |
| | **Hydrogen bond<br>1:THR308(HG1-O)1:THR308** | **1.947** |
| | **Hydrogen bond<br>1:THR309(H-O)1:ASP300** | **3.02** |
| KIR2.1-WT_OPEN | Hydrogen bond<br>2:THR308(HG1-O)2:THR308 | 2.203 |
| | Hydrogen bond<br>2:GLU299(OE1-H) 2:GLY300 | 2.003 |
| | Hydrogen bond<br>2:THR308(CA-SD)2:MET301 | 4.019 |
| G300D_CLOSED | Hydrogen bond<br>2:THR308(HG1-O)2:THR308 | 2.095 |

| | | |
|---|---|---|
| | Hydrogen bond 2:THR308(HG1-OD2)2:ASP300 | 1.646 |
| | Hydrogen bond 2:THR308(CA-SD)2:MET301 | 3.945 |
| KIR2.1-WT_OPEN | Hydrogen bond 3:THR308(HG1-O)3:THR308 | 2.008 |
| | Hydrogen bond 3:THR309(HG1-O) 3:GLY300 | 1.875 |
| G300D_CLOSED | Hydrogen bond 3:THR308(HG1-O)3:THR308 | 1.930 |
| | Hydrogen bond 3:THR309(HG1-OD2) 3:ASP300 | 1.637 |
| KIR2.1-WT_OPEN | Hydrogen bond 4:THR308(HG1-O)4:THR308 | 1.989 |
| | Hydrogen bond 4:GLU224(OE2-CA) 4:GLU224 | 2.995 |
| G300D_CLOSED | Hydrogen bond 4:THR308(HG1-O)4:THR308 | 2.000 |
| | Hydrogen bond 4:THR308(HG1-O)4:MET307 | 2.230 |
| | Hydrogen bond 4:THR308(CB-OD2)4:ASP300 | 3.493 |
| | Hydrogen bond 4:GLU224(OE2-CA) 4:GLU224 | 2.978 |

*****(Figure 3-5) in the main manuscript are highlighted in **bold** in the Table 1.

1. Fernandes, C.A. et al. Cryo–electron microscopy unveils unique structural features of the human Kir2. 1 channel. *Science Advances* **8**, eabq8489 (2022).
